# Impaired adipose anabolism in pancreatic cancer cachexia is reversed by HuR inhibition

**DOI:** 10.1101/2024.12.27.630549

**Authors:** Paige C. Arneson-Wissink, Katherine Pelz, Beth Worley, Heike Mendez, Peter Pham, Parham Diba, Peter R. Levasseur, Grace McCarthy, Alex Chitsazan, Jonathan R. Brody, Aaron J. Grossberg

**Author notes:** Corresponding Author: Aaron J. Grossberg, 3181 SW Sam Jackson Park Rd Mail Code L481, Oregon Health and Science University, Portland, OR, 97239.

## Abstract

**Background:** Cachexia is defined by chronic loss of fat and muscle, is a frequent complication of pancreatic ductal adenocarcinoma (PDAC), and negatively impacts patient outcomes. Nutritional supplementation cannot fully reverse tissue wasting, and the mechanisms underlying this phenotype are unclear. This work aims to define the relative contributions of catabolism and anabolism to adipose wasting in PDAC-bearing mice. Human antigen R (HuR) is an RNA-binding protein recently shown to suppress adipogenesis. We hypothesize that fat wasting results from a loss of adipose anabolism driven by increased HuR activity in adipocytes of PDAC-bearing mice.

**Methods:** Adult C57BL/6J mice received orthotopic PDAC cell (*Kras^G12D^; p53^R172H/+^; Pdx1-cre*) (PDAC) or PBS (sham) injections. Mice exhibiting moderate cachexia (9 days after injection) were fasted for 24h, or fasted 24h and refed 24h before euthanasia. A separate cohort of PDAC mice were treated with an established HuR inhibitor (KH-3, 100 mg/kg) and subjected to the fast/refeed paradigm. We analyzed body mass, gross fat pad mass, and adipose tissue mRNA expression. We quantified lipolytic rate as the normalized quantity of glycerol released from 3T3-L1 adipocytes *in vitro*, and gonadal fat pads (gWAT) *ex vivo*.

**Results:** 3T3-L1 adipocytes treated with PDAC cell conditioned media (CM) had lower expression of lipolysis and lipogenesis genes than control cells, and did not display elevated lipolysis as measured by liberated glycerol. PDAC gWAT cultured *ex vivo* displayed decreased lipolysis compared to sham gWAT (-54.7%). PDAC and sham mice lost equivalent fat mass after a 24h fast, however, PDAC mice could not restore inguinal fat pads (iWAT) (-40.5%) or gWAT (-31.8%) mass after refeeding. RNAseq revealed 572 differentially expressed genes in gWAT from PDAC compared to sham mice. Downregulated genes (n=126) were associated with adipogenesis (adj p=0.05), and expression of adipogenesis master regulators *Pparg* and *Cebpa* were reduced in gWAT from PDAC mice. Immunohistochemistry revealed increased HuR staining in gWAT (+74.9%) and iWAT (+41.2%) from PDAC mice. Inhibiting HuR binding restored lipogenesis in refed animals with a concomitant increase in iWAT mass (+131.7%).

**Conclusions:** Our work highlights deficient adipose anabolism as a driver of reduced lipid content in 3T3-L1 adipocytes treated with PDAC conditioned media and PDAC mice. The small molecule KH-3, which disrupts HuR binding, restored adipose anabolism in PDAC mice. This highlights HuR as a potentially targetable regulatory node for adipose anabolism in cancer cachexia.

## INTRODUCTION

Cancer-associated cachexia is a wasting condition characterized by systemic inflammation, progressive weight loss, and atrophy of white adipose tissue (WAT) and skeletal muscle [1, S1-2]. In addition to physical deterioration, individuals with cachexia also exhibit fatigue, anorexia, and cognitive decline [1, S1-3], which contribute significantly to reductions in quality of life, ability to tolerate chemotherapy or surgery, and patient mortality [2, 3, S4-6]. Cachexia is estimated to be the direct cause of death in 20-30% of cancer patients [2, S4], and among all malignancies, pancreatic ductal adenocarcinoma (PDAC) is the most highly associated with cachexia, with an estimated 83% of patients suffering from this condition [4, S7-8]. Despite much of cachexia research focusing on improving skeletal muscle mass, a retrospective study of patients with PDAC revealed that fat loss alone is associated with equally poor outcomes as combined muscle and fat mass loss [5]. Additionally, current cachexia clinical trials center around weight maintenance or gain as a primary endpoint. This highlights the importance of understanding the drivers of adipose tissue loss in cachexia.

Cachexia arises when energy catabolism exceeds anabolism, leading to unsustainable levels of fat mobilization and muscle depletion. Multiple factors are known to enhance catabolism, including decreased secretion of anabolic hormones, and altered metabolism of protein, carbohydrate, and lipid substrates [6]. Current work suggests that inflammation drives metabolic abnormalities in cachexia [7, S7]. Pro-inflammatory cytokine activity increases during cancer progression [8, 9] and systemic inflammation can contribute to wasting by inducing hypercatabolism in muscle and adipose tissue [6, 10–12]. Existing literature highlights elevated rates of lipolysis and adipose browning as the primary forces underlying adipose tissue wasting [13, 14]. Browning, in particular, appears to exert a double effect in cachexia by both reducing lipid stores and increasing energy expenditure [14]. However, impaired anabolic processes, like adipogenesis and lipogenesis, also contribute to adipose tissue loss [15]. Targeting peroxisome proliferator-activated receptor gamma (PPARG), a transcriptional control point of adipogenesis, with the agonist rosiglitazone was sufficient to improve fat and muscle mass retention in mice with lung cancer [16]. Tumor-derived factors are also capable of impairing adipogenesis in cultured 3T3L-1 adipocytes [17, 18].

RNA-binding proteins (RBPs) are essential in governing biogenesis, stabilization, translation, and decay of mRNA transcripts [19, 20]. In adipose tissue, several RBPs are documented to regulate alternative splicing [21–23], control of key adipogenic transcription factors [24], and the translation efficiency of proteins involved in browning [25]. Until recently, the function of most RBPs in adipocytes was largely unexplored. The RBP human antigen R (HuR), encoded by the embryonic lethal abnormal vision-like 1 (*ELAVL1*) gene, regulates the expression of genes involved in inflammation, stress response, and apoptosis [26]. Although HuR likely exerts complex effects in the context of both PDAC and cachexia, we selected HuR as a target of interest because recent work showed that it is a repressor of adipogenesis in both white and brown adipose tissues [27]. Our goal was to elucidate the relative contributions of enhanced catabolism and impaired anabolism on fat wasting by investigating adipose tissue response to different nutritional contexts and HuR inhibition in cachectic mice.

## METHODS

### Cell culture

#### KPC PDAC cells

KPC cells expressing pancreas-specific conditional alleles (*Kras^G12D^; p53^R172H/+^; Pdx1-cre*) [28] were stored in liquid nitrogen until use and then maintained in RMPI 1640 supplemented with 10% FBS, 1mM sodium pyruvate, and 50 U/mL penicillin/streptomycin (Gibco, Gaithersburg, MD) at 37C and 5% CO2. Conditioned media (CM) was collected from confluent KPC cells grown in DMEM, 1% pen/strep (to accommodate later culturing of 3T3-L1 cells) after 24 hours incubation, centrifuged at 1,200 g for 10 min, filtered with a 0.2-μm syringe filter, and used immediately or stored at −80°C.

#### 3T3-L1 adipocytes

3T3-L1 adipocytes (ATCC CL-173) were purchased from ATCC and stored in liquid nitrogen until use. Cells were cultured in preadipocyte expansion media (DMEM, 10% bovine calf serum, 1% pen/strep). Then, cells were plated for differentiation at a density of 2x10^5^ cells/6-well dish, or 6.7 x10^3^ cells/96well dish and maintained in preadipocyte expansion media for 2 days after reaching 100% confluency. Media was changed to adipocyte differentiation media (DMEM, 10% FBS, 1% pen/strep, 1 uM dexamethasone, 0.5 mM IBMX, 1 ug/mL bovine insulin) for 2 days before switching to adipocyte maintenance media (DMEM, 10% FBS, 1% pen/strep, 1 ug/mL bovine insulin) for the duration of the experiment. Cells were cultured for up to 6 days after start of maintenance media to reach complete differentiation. For Oil Red O staining, CM was added in place of adipocyte maintenance media and was supplemented with FBS and insulin for 6 days. For lipolysis assay (media glycerol, qPCR, and western blot), cells were completely differentiated and then changed to control media, KPC conditioned media without insulin or FBS. Isoproterenol (10 uM) was added to media as a positive control. Lipolysis end points were collected at 1, 3, and 24 hours after media change.

#### Oil red O staining

Diluted Oil Red O stock solution was prepared by mixing concentrated Oil Red O (0.5% Oil Red O in 100% isopropyl alcohol) 6:4 with deionized water. This solution was allowed to stand for 10 minutes at 4C and then filtered immediately before use through a 0.22 um filter. 3T3-L1 cells were washed with PBS, then incubated in 4% paraformaldehyde for 30 minutes at room temperature, then washed with deionized water twice. Cells were incubated in 60% isopropyl alcohol for 5 minutes, then treated with diluted Oil Red O solution for 15 minutes. Cells were washed 5 times with deionized water, then red signal was quantified at 490 nm in a plate reader (BioTek).

#### Glycerol quantification

3T3-L1 adipocyte media was collected and run at a 1:2 dilution, and mouse plasma samples were run undiluted for glycerol quantification according to the manufacturer’s instructions (Abcam ab65337). Fresh media and KPC CM alone (no 3T3-L1 exposure) were below the detection limit of the assay.

#### NEFA quantification

Mouse plasma was run undiluted according to the manufacturer’s instructions (Wako Diagnostics NEFA-HR series). mEq/L are the units specified by the assay manufacturer and what is standard reporting in clinical values. mEq/L = mM * |valence charge| . Because NEFAs have a valence charge of -1, mEq/L is equivalent to mM.

#### Mouse studies

Wildtype C57BL/6J mice (JAX catalog number 000664) were purchased from The Jackson Laboratory (Bar Harbor, ME) and maintained in standard rodent housing at 26°C with 12h light/12h dark cycles. Sex is defined in individual figure legends. Animals used for experimentation were 12-15 weeks of age. Mice were individually housed for acclimation for 7 days prior to tumor implantation and provided *ad libitum* access to water and food (Rodent diet 5001; Purina Mills, St. Louis, MO, USA). Food intake was measured daily. Tumor-bearing animals were euthanized by cardiac puncture under deep isoflurane anesthesia. Studies were conducted in accordance with the U.S. National Institutes of Health Guide for the Care and Use of Laboratory Animals and approved by the Institutional Animal Care and Use Committee of Oregon Health & Science University.

#### Study-specific manipulations

For fasting studies, mice were transferred to clean cages without food for 24 hours prior to euthanasia or refeeding. Pair-feeding was used on the study days indicated in the figure legends by feeding sham mice the average of the food consumed by PDAC mice the day prior. PDAC mice treated with the HuR inhibitor KH-3, were injected intraperitoneally (100 mg/kg) at 6-, 8-, and 10-days post-implantation.

#### Orthotopic PDAC implantation

Wildtype C57BL/6J mice aged 12-15 weeks received orthotopic PDAC tumor injections (*Kras^G12D^; p53^R172H/+^; Pdx1-cre*) or sham injections. PDAC mice were injected with 1x10^6^ KPC cells into the tail of the pancreas parenchyma in a volume of 23ul while sham animals were treated with an equal volume of PBS. Animals were euthanized 10-11 days after tumor implantation. N and sex are defined on a per-study basis in the figure legends.

#### Echo magnetic resonance imaging body composition

Lean mass, fat mass, total body water, and free water were measured using whole-body magnetic resonance imaging (MRI) (EchoMRI, Houston, TX). Measurements were taken pre-implantation (baseline), pre-fasting and at euthanasia in tumor and sham groups to assess body composition.

#### Tissue collection and histology

Tissues collected at necropsy were weighed and flash frozen in liquid nitrogen prior to storage at -80C. Tissues for HuR histology were fixed with 4% paraformaldehyde overnight and then transferred to 70% ethanol. Tissues were paraffin-embedded, sectioned, incubated with anti-HuR (1:300 #sc-5261, Santa Cruz) and stained using horseradish peroxidase-conjugated secondary antibody and incubation in 3,3’-diaminobenzidine by the Histopathology Shared Resource Core at OHSU. Whole tissue sections were scanned by the Advanced Light Microscopy Core at OHSU. ZEN Digital Imaging for Light Microscopy (RRID:SCR_013672) was then used to obtain at least five 10x images per tissue. ImageJ software was used to quantify the percent staining for each section (56). All staining was quantified by running color deconvolution on 10x images, applying a standard intensity threshold on the corresponding images, and measuring the percent area covered by the staining.

#### Ex vivo lipolysis

*Ex vivo* lipolysis assays of adipose explants were performed as previously described [29]. Briefly, gonadal white adipose tissue (gWAT) tissue was collected from ad-lib fed sham and PDAC mice 10 days after tumor implantation. Tissues were cut into approximately 100 mg samples, minced, then incubated in phenol red-free DMEM containing 2% fatty acid free bovine serum albumin at 37°C for 1 hour. Media was collected and snap-frozen. The tissue was transferred to a new dish containing media only or media with 10 uM isoproterenol and incubated for 2 hours at 37C. Media was collected and snap-frozen. Media aliquots were thawed and analyzed for glycerol using Sigma Glycerol Assay Kit (MAK117) according to manufacturer’s directions.

#### Quantitative qRT-PCR

Total RNA was extracted from cell pellets or tissues with the E.Z.N.A. Total RNA Kit II (Omega Bio-Tek Inc., Norcross, GA). cDNA was transcribed with the High-Capacity cDNA Reverse Transcription Kit (Applied Biosystems, Waltham, MA). Quantitative real-time polymerase chain reaction (qPCR) was run on the ABI 7300 (Applied Biosystems) using TaqMan Fast Advanced PCR Master Mix (Applied Biosystems) or SYBR Green Master Mix (Applied Biosystems). The relative expression was calculated using the ΔΔCt method with gene expression relative to beta actin or 18S.

#### RNA sequencing

RNA sequencing was performed on total RNA isolated as described above. RNA libraries were prepared and sequenced using the Illumina Nova Seq and HiSeq platform according to the Illumina Tru-Seq protocol (Novogene, Sacramento, CA). Detailed analysis methods are included in the supplement.

#### Western Blot

3T3-L1 protein was collected by scraping cells into lysis buffer followed by brief sonication. Adipose tissue protein was collected using the Minute Total Protein Extraction Kit (Invent Biotechnologies Inc AT-022) and the manufacturer’s protocol. We used BCA assay to determine protein concentration. Approximately twenty micrograms of protein was loaded in each lane and run on 4–12% Bis-Tris NuPAGE gel (Invitrogen). Gels were transferred to polyvinylidene difluoride (PVDF) membranes (Millipore) and blocked with 5% BSA for 1 h.

Membranes were incubated with primary antibodies overnight at 4°C with gentle agitation. Blots were then washed with Tris-buffered saline with Tween 20 (TBST) and incubated in secondary antibodies for 1 h prior to imaging (LI-COR Odyssey Imaging System). Ladders used were iBright Pre-stained Protein Ladder (Invitrogen)(HuR blot), and Chameleon 700 Pre-stained Protein Ladder (LI-COR)(all other blots). Primary antibodies: Vinculin (Santa Cruz sc-73614), HuR (Santa Cruz sc-5261), ATGL (Cell Signaling 2138), HSL (Cell Signaling 4107), Phospho-HSL (Ser660) (Cell Signaling 45804), and Phospho-HSL (Ser563) (Cell Signaling 4139). Secondary antibodies: Mouse IgG and Rabbit IgG Dylight 680 and 800 (Cell Signaling), goat anti-mouse IgG Alexa Fluor Plus 800 (Invitrogen A32730), and goat anti-mouse IgG Alexa Fluor Plus 647 (Invitrogen A32728).

#### Statistical Analysis

Specific statistical tests and sample size for each study is indicated in the figure legends. Error bars in figures show SEM. Statistical analyses were performed using GraphPad Prism (version 9; GraphPad Software Inc), and graphs were built using GraphPad Prism (GraphPad Software Inc) statistical analysis software or R studio (4.2.3) using ggplot (3.5.1). P values are 2 sided with values less than 0.05 regarded as statistically significant.

#### Data Availability

Further information and resources, including KPC cells and raw data will be shared upon reasonable request to Aaron J. Grossberg.

## RESULTS

### Adipocytes suppress lipogenesis in response to PDAC-derived factors

Given a large body of literature suggesting that PDAC cachexia is partly driven by increased lipolysis in the adipose tissue, we tested the effect of PDAC cell-conditioned media (KPC CM) on lipolysis *in vitro* [13, 14, 30]. We treated differentiated 3T3-L1 cells with control maintenance media, or with KPC CM for 6 days. At this time, we observed decreased lipid droplet accumulation by brightfield microscopy, which was confirmed with Oil Red O (**Figure 1A-B**). Media glycerol levels, a measure of lipolysis, were not elevated in KPC CM-treated cells (**Figure 1C**, **S1A**). Correspondingly, protein markers of lipolysis, including activated (phosphorylated) hormone-sensitive lipase (HSL) and adipose triglyceride lipase (ATGL) were not elevated in KPC CM-treated cells (**Figure 1D-E, S1B-E, S2**) and mRNA levels of lipolysis enzymes *Atgl* and hormone-sensitive lipase (lipase e, *Lipe*) were significantly decreased in KPC CM-treated 3T3-L1 cells (**Figure 1F**). To explain the apparent decrease in lipid accumulation without increased lipolysis, we measured the expression of genes associated with lipogenesis: diacylglycerol O-acyltransferase 1 and 2 (*Dgat1 and Dgat 2),* fatty acid synthase (*Fasn*), lipoprotein lipase (*Lpl*), nuclear receptor subfamily 1 group H member 3 (*Nr1h3*), stearoyl-CoA Desaturase 1 (*Scd1*), solute carrier family 2 member 4 (*Slc2a4*), and sterol regulatory element-binding protein 1c (*Srebp1c*). All of these genes were suppressed after 24 hour KPC CM treatment, indicating that the decreased lipid content in 3T3-L1 cells was due to impaired anabolic activity rather than increased catabolic activity (**Figure 1F**).

**Figure 1:**
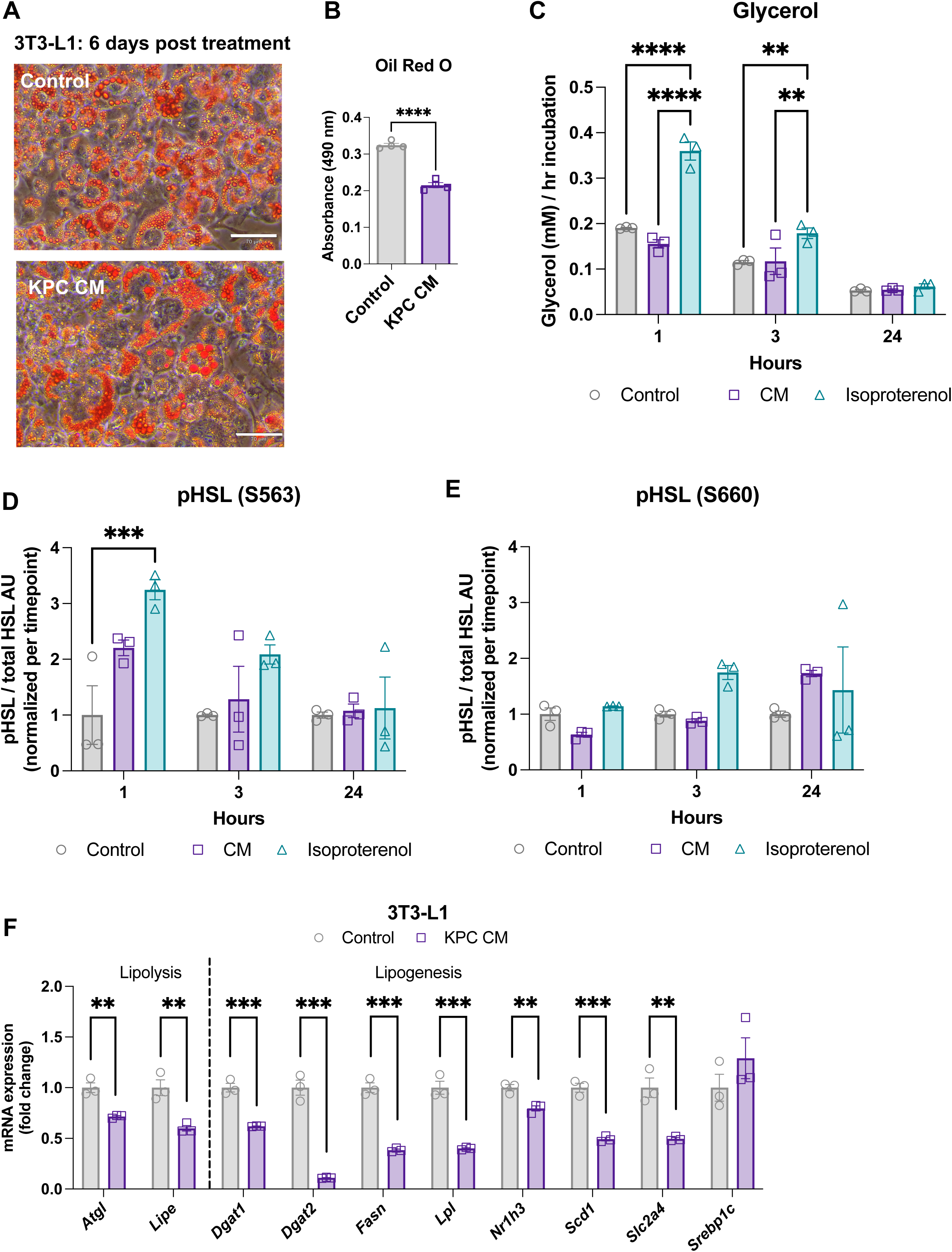
Adipocytes suppress lipogenesis in response to pancreatic ductal adenocarcinoma-conditioned media. **(A**) Representative brightfield images of differentiated 3T3-L1 cells treated with control media (top) or KPC CM (bottom) for **6** days. Scale bars represent 70 um. **(B)** Quantification of Oil Red O staining. N = 4 wells per condition. (**C-E**) 3T3-L1 cells were fully differentiated and then treated with control, KPC-conditioned, or isoproterenol (10 uM) media for 1, 3, and 24 hours in the absence of FBS and insulin. N = 3 wells per timepoint/condition. (**C**) Media glycerol levels, normalized by hours of collection . Conditioned media and control media alone did not contain glycerol above background (assay buffer only) levels. (**D**) Serine 563 phosphorylated HSL protein levels normalized to total HSL protein levels. (**E**) Serine 660 phosphorylated HSL protein levels normalized to total HSL protein levels. (**F**) mRNA expression of lipolysis and lipogenesis genes. N= 3 wells per condition, normalized to 18S expression. Panels B and F tested with t-test. Panels C-E tested with two-way ANOVA with Tukey correction for multiple comparisons. p<0.05, **p<0.01, ***p<0.001, and ****p<0.0001.

### PDAC is associated with decreased fat pad mass *in vivo*

We first sought to assess the effects of pancreatic cancer on metabolism by characterizing tissue physiology in a murine orthotopic PDAC model. 12-week-old C57BL/6J mice with PDAC or sham implantations were fed *ad libitum* with or without a 24h fast once mice reached a moderate cachexia burden (9 days after tumor implantation). Body mass maintained stable among all groups prior to fasting (**Figure 2A**), whereas PDAC mice exhibited a small reduction in food intake (**Figure 2B**) compared to sham controls. Fasting caused a significant reduction in tumor mass compared to *ad libitum*-fed mice **(Figure 2C).** EchoMRI body composition analysis of fat mass demonstrated significant decreases in overall adiposity in PDAC animals from baseline to pre-fast due to cachexia progression and reduction in food intake (**Figure 2D**). As expected, between pre- and post-fast, both PDAC and sham animals lost significant overall adiposity (**Figure 2E**). Terminal fat pad masses (inguinal, iWAT, and gonadal, gWAT) were significantly decreased in both ad lib and fasted conditions as compared to sham mice (**Figure 2F**). PDAC mice were also vulnerable to skeletal muscle mass loss as measured by echoMRI and gastrocnemius muscle tissue mass (**Figure S3**).

**Figure 2:**
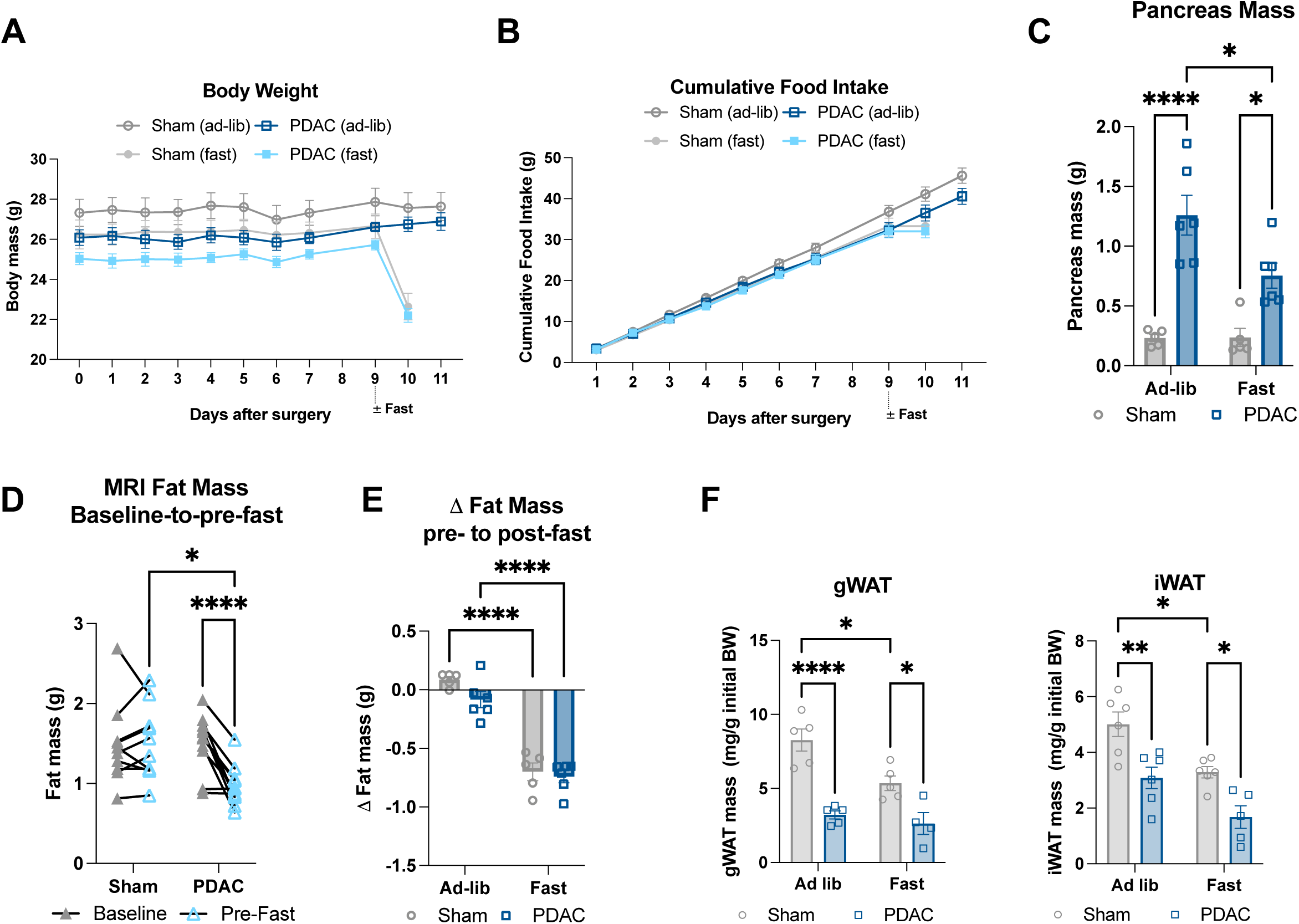
Pancreatic ductal adenocarcinoma is associated with decreased fat pad mass *in vivo*. (**A-E**)Wildtype C57BL/6J mice with PDAC or sham implantations were fed *ad libitum* or fasted 16 h at mid-cachexia (9 days after injection). Animals were euthanized 10 or 11 days after tumor implantation following a 16 h fast. N=5 sham, 6 PDAC male mice per feeding condition. **(A)** Daily body mass. Statistically tested with 3-way ANOVA p<0.0001 for time, time x fast; p=0.0073 for fast; p=0.0097 for time x tumor status; p=0.0167 for tumor status. (**B**) Cumulative food intake. Statistically tested with 3-way ANOVA p<0.0001 for time, time x fast; p=0.0035 for time x tumor status. (**C**) Terminal pancreas/tumor mass. (**D-E**) Body composition changes in total adiposity were characterized from baseline (pre-tumor implantation) to pre-fast between sham and PDAC mice (**C**), and from pre- to post-fast (**D**). (**F**) Terminal gWAT and iWAT mass following 24 h fast 14 days after injection. N=5 male mice per *ad lib* group, 5 male mice sham fast, 3 male mice PDAC fast. 2x2 analyses were statistically tested with two-way ANOVA or mixed effects model with Tukey multiple comparisons. *p<0.05, **p<0.01, and ****p<0.0001.

### PDAC impairs lipolysis and adipogenesis

Based on our findings *in vitro*, we hypothesized that PDAC mice would not exhibit increased lipolysis, which is characteristic of other cachexia models [13, 30]. To assess products of lipolysis in circulation, we measured plasma glycerol and non-esterified fatty acids (NEFAs). Plasma glycerol levels were significantly lower in fasted PDAC mice than in fasted sham mice (**Figure 3A**), while there was no significant difference in plasma NEFA levels between fasted groups (**Figure 3B**). To directly measure the rate of adipose tissue catabolism, we performed an *ex vivo* lipolysis study, using gonadal white adipose tissue (gWAT) collected from sham and PDAC mice. We assessed fat pads both in the presence and absence of the beta-adrenergic agonist isoproterenol (10 uM) to determine baseline and stimulated lipolysis [29]. Baseline and isoproterenol-stimulated glycerol release were significantly decreased in PDAC gWAT (**Figure 3C**), demonstrating that PDAC suppresses, rather than enhances, lipolytic rate and capacity. To further characterize adipose tissue in PDAC mice, we assessed mRNA expression of genes regulating lipolysis and browning in adipose tissue. PDAC mice had suppressed protein and mRNA expression of ATGL and suppressed activation (phosphorylation) of HSL in the gWAT (**Figure 3D,F, S4**). PDAC mice also showed suppressed iWAT and gWAT *Lipe* expression under fasted conditions (**Figure 3D-E**). We observed suppressed browning-associated gene expression in gWAT, including the genes peroxisome proliferator-activated receptor gamma coactivator 1 alpha (*Pgc1a),* protein domain containing 16 (*Prdm16*), and cell death inducing DFFA like effector a (*Cidea*) in PDAC mice in both *ad libitum* fed and fasted conditions. Uncoupling protein 1 (*Ucp1*), which is also associated with browning, was not significantly increased in gWAT or iWAT from PDAC mice (**Figure 3D-E**). Therefore, neither enhanced lipolysis nor browning are significant contributors to adipose loss in this model.

**Figure 3:**
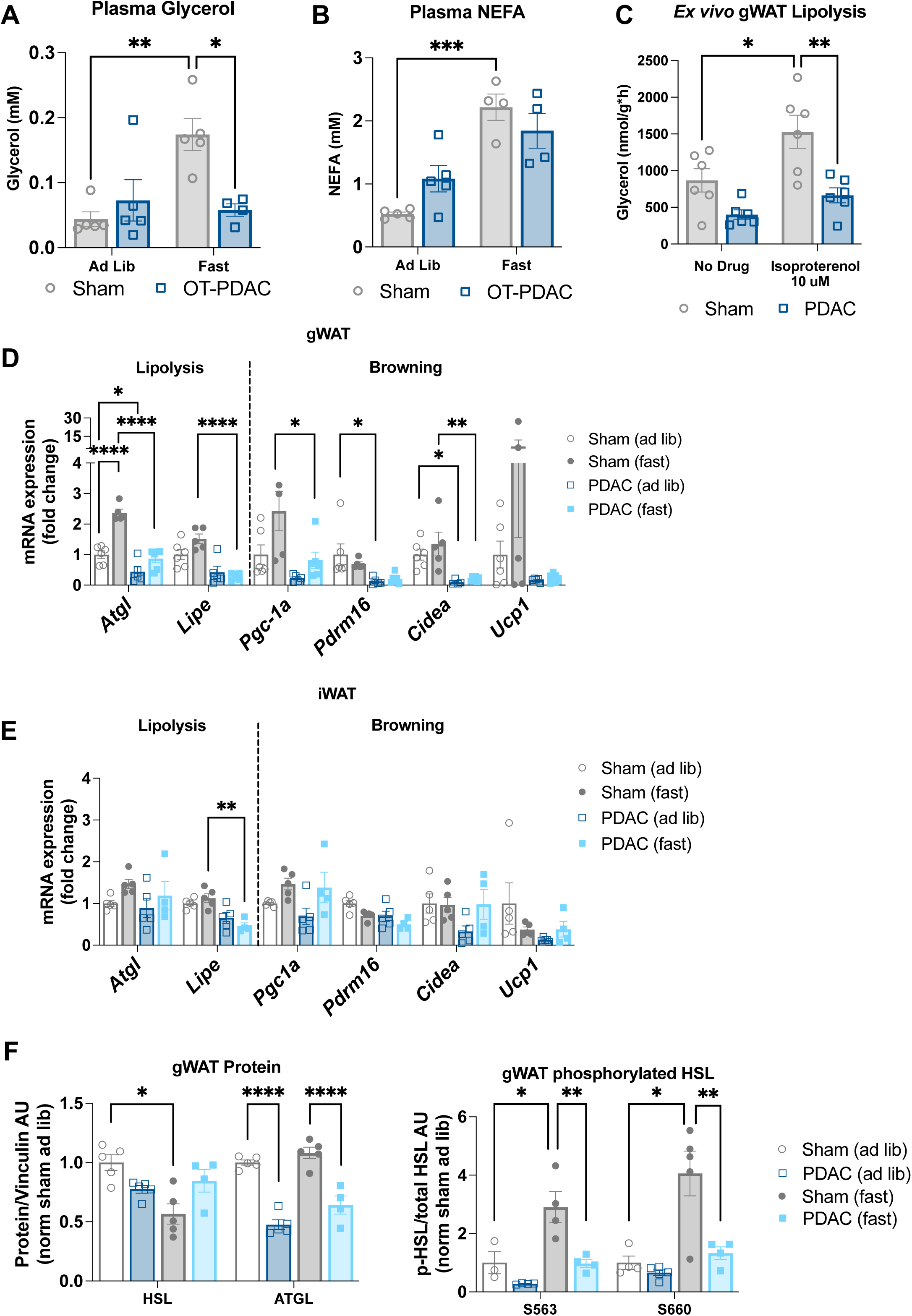
Pancreatic ductal adenocarcinoma impairs lipolysis and adipogenesis *in vivo*. **(A)** Measurement of lipolysis (glycerol release) in explants derived from gWAT collected 10 days after tumor implantation. Lipolysis was induced by the β-adrenergic agonist isoproterenol (10 µM). N=6 male mice per group. Statistically tested with two-way ANOVA with uncorrected Fisher’s LSD test to compare groups. (**D**) mRNA levels of lipolysis and browning genes in gWAT collected 10 days post-implantation from 24 h-fasted and *ad libitum*-fed mice, normalized to 18S expression. N=5 male mice sham/fasted; 3 male, 3 female mice sham/ad lib; 6 male mice PDAC/fasted; 3 male, 2 female mice PDAC/ad lib. (**E**) mRNA expression of lipolysis and browning genes in iWAT collected 11 days post-implantation from *ad libitum*-fed mice, normalized to beta actin expression. N=4 male mice per group. (**F**) Western blot analysis of PDAC and Sham gWAT protein for HSL (83 kDa) and ATGL (55 kDa) (left) and Serine 563- and Serine 660-phosphorylated HSL protein levels normalized to total HSL protein levels (right). N=5 male mice per group. Protein normalized to vinculin (124 kDa). 2x2 data were statistically tested with two-way ANOVA with Tukey multiple comparisons comparing tumor experimental groups to sham controls. Pairwise comparisons were made with unpaired t-test. *p<0.05, **p<0.01, ***p<0.001, and ****p<0.0001.

### PDAC downregulates pathways associated with adipogenesis

To gain further insights into PDAC metabolism from global gene expression analysis, we performed bulk RNA sequencing (RNAseq) on gWAT from sham and PDAC animals. . In gWAT, we identified a total of 572 differentially expressed genes (DEGs) in PDAC versus sham mice, of which 446 were enriched, and 126 were depleted (**Figure 4A, S5**). Pathway analysis referencing the molecular signatures database (MSigDB) hallmark gene set collection revealed increased expression of gene sets associated with cell cycle control (e.g., E2F targets and G2M checkpoints) and inflammation (e.g. TNFa and JAK/STAT3 signaling) and decreased expression of genes linked to adipogenesis, oxidative phosphorylation, and fatty acid metabolism (**Figure 4B**) [31]. Because inhibition of adipogenesis could provide an alternative mechanism for adipose wasting, we plotted all significant DEGs in the adipogenesis gene set and observed nearly universal depletion of these transcripts in PDAC mice (**Figure 4C**). We then validated our RNAseq data by performing qPCR on selected adipogenesis-associated DEGs, revealing consistent downregulation of both adipogenesis and lipogenesis genes (**Figure 4D**). We then repeated qPCR in iWAT to determine whether subcutaneous adipose exhibited the same transcriptomic changes. Transcriptional changes in iWAT were less dramatic than in gWAT and did not reach significance, with the exception of *Lpl*, which was significantly decreased in PDAC tissue (**Figure 4E**). Together, these results indicate that suppressed transcriptomic programs associated with adipogenesis could account for decreased adipose tissue during cachexia, in the absence of elevated lipolysis.

**Figure 4:**
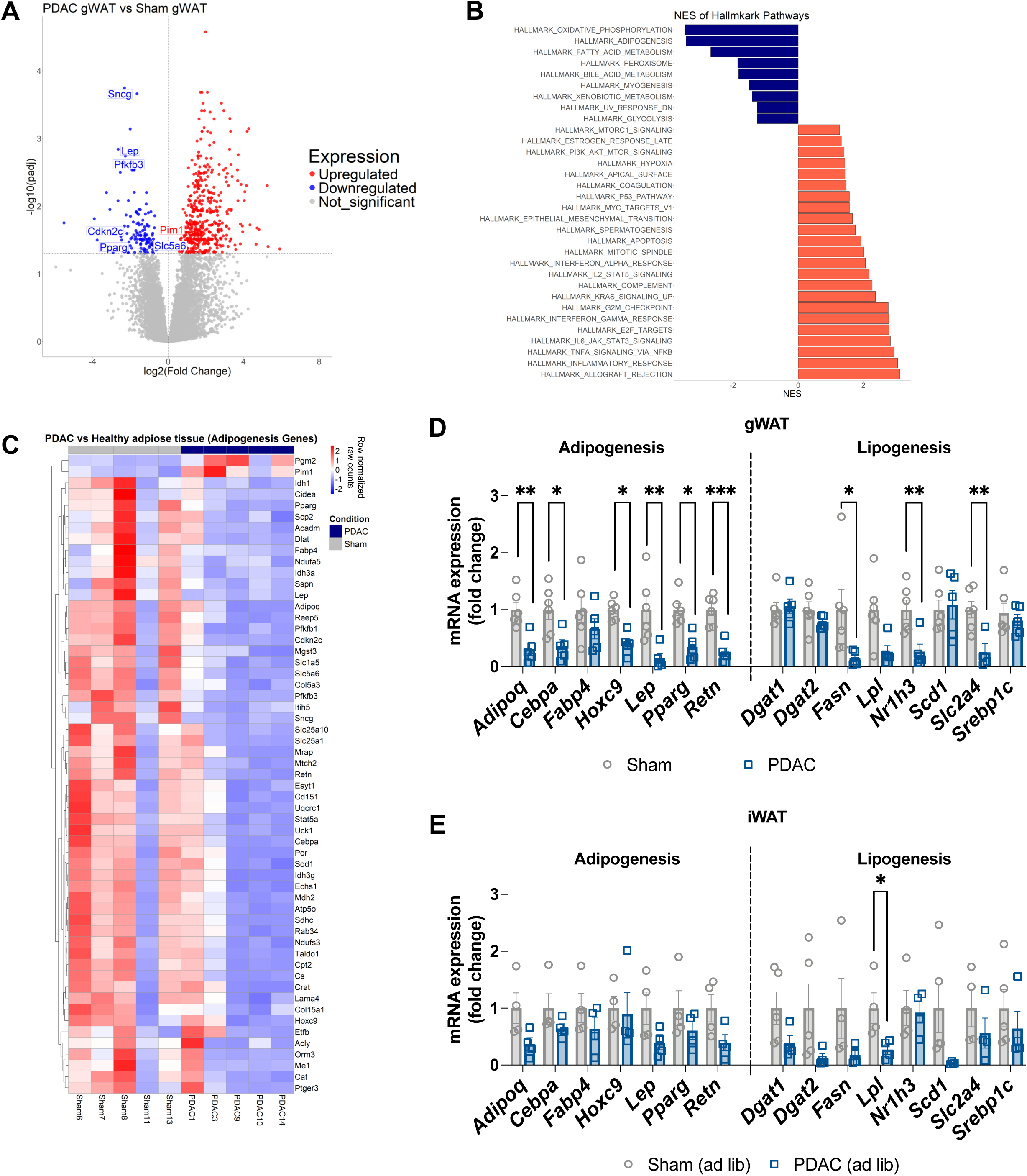
PDAC downregulates pathways associated with adipogenesis. RNAseq on gWAT from *ad libitum*-fed PDAC and sham mice collected 14 days post tumor implantation. N=2 female, 3 male PDAC; 3 female, 2 male sham. **(A)** Volcano plot of differentially expressed genes in gWAT. Blue dots represent significantly downregulated genes (log2 fold change < -1 and adj p<0.05). while red dots represent significantly upregulated genes (log2 fold change >1 and adj p<0.05). Grey dots are insignificant. Labels denote significant differentially expressed genes from the Hallmark Adipogenesis pathway. **(B)** Broad Institute GSEA showing significantly enriched pathways in PDAC gWAT relative to sham gWAT (pathway p<0.05). Blue is decreased expression, red is increased expression. **(C)** Heat map of Hallmark Adipogenesis pathways constituents in PDAC and Sham gWAT. Scale represents row-normalized raw counts. **(D)** qPCR validation of adipogenesis and lipogenesis genes in in gWAT collected 10 days post-implantation from 24 h-fasted and *ad libitum*-fed mice. N=5 male mice sham/fasted; 3 male, 3 female mice sham/ad lib; 6 male mice PDAC/fasted; 3 male, 2 female mice PDAC/ad lib. Statistically tested with two-way ANOVA with Tukey multiple comparisons comparing tumor experimental groups to sham controls. **(E)** qPCR validation of adipogenesis and lipogenesis genes in iWAT from ad lib fed mice collected 11 days post implantation. N=4 male mice per group. Statistically tested with unpaired t-test. *p<0.05, **p<0.01, ***p<0.001, and ****p<0.0001.

### Adipose tissue anabolism is impaired in orthotopic PDAC mice after refeeding

Based on our results demonstrating downregulation of adipogenic and lipogenic genes in PDAC mice, we next wanted to confirm whether adipose tissue anabolism is functionally impaired in mice implanted with orthotopic PDAC. To do this, wildtype C57BL/6J mice with PDAC or sham implantations were fed *ad libitum* for 9 days, an established timepoint of active cachexia [32], fasted 24h, and then terminated or allowed to re-feed for 24 h. To control for differences in caloric intake, we used a pair-fed refeeding scheme in which sham mice were refed with the average food consumption of the refed PDAC group. Challenging mice with a 24h fast depletes adipose mass in both PDAC and sham mice (**Figure 2D**), enabling us to assess anabolic restoration of fat mass during a 24 h refeeding period. We measured cumulative food intake and daily body mass for the duration of the study (**Figure 5A-B**). 24h fasting followed by 24h refeeding did not impact pancreas tumor mass (**Figure S6**). There were no differences in iWAT or gWAT mass in fasted PDAC versus sham mice (**Figure 5C-D**). However, while sham mice regain significant amounts of gWAT and iWAT mass after refeeding, gWAT and iWAT masses in PDAC mice remain equivalent before and after refeeding (**Figure 5C-D**). Refeeding sham mice caused increased gWAT expression of lipogenic *Fasn* and adipogenic genes *Lep* and *Retn*, while these genes remained suppressed in PDAC gWAT (**Figure 5E**). Refed sham iWAT showed increased *Fasn,* but not adipogenic genes, while PDAC iWAT had suppressed expression of both adipogenic and lipogenic genes (**Figure 5F**). These results confirm that downregulation in pathways mediating adipogenesis are indeed accompanied by impaired adipose tissue anabolism in orthotopic PDAC mice after refeeding.

**Figure 5:**
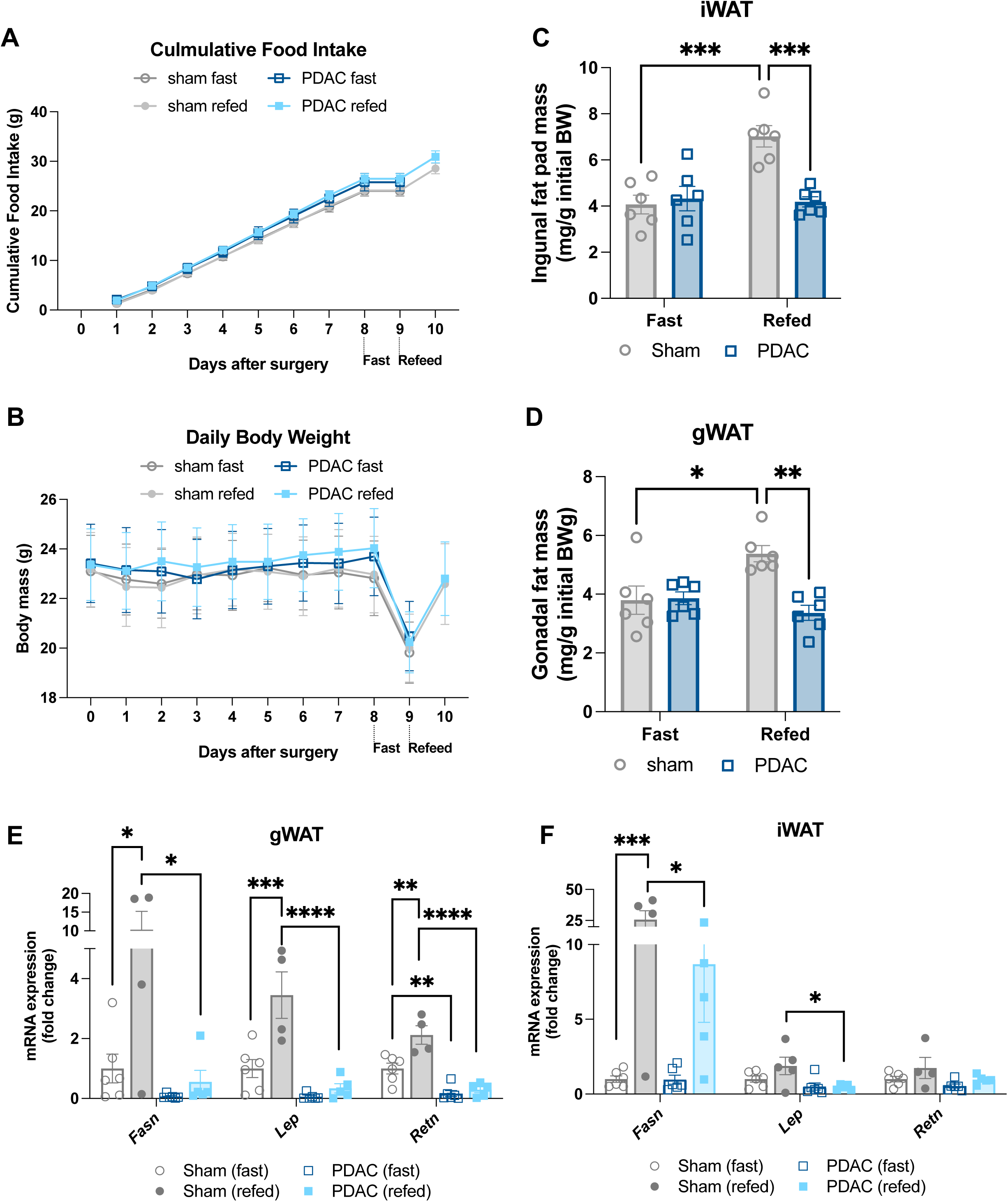
Adipose tissue anabolism is impaired in orthotopic PDAC mice after refeeding. Wildtype C57BL/6J mice with PDAC were fed *ad libitum* for 9 days then fasted 24 h with or without a 24 h refeeding period. Sham mice were pair-fed daily to the average food intake of the PDAC group from 2 days post-implantation until study end. Animals were euthanized 10 or 11 days after tumor implantation following the 24 h fast (10 d) or 24 h refeed (11 d). N=3 male and 3 female mice per group. **(A)** Cumulative food intake. Statistically tested with 3-way ANOVA. P<0.0001 for time, p=0.0344 for time x tumor status. **(B)** Daily body mass. Statistically tested with 3-way ANOVA. P<0.0001 for time, p=0.002 for time x tumor status. iWAT (**C**) and gWAT (**D**) mass normalized to initial body mass. mRNA expression of anabolic genes in gWAT (**E**) and iWAT (**F**), normalized to beta actin expression. Data represented in C-F were statistically tested with two-way ANOVA with Tukey multiple comparisons comparing PDAC groups to sham controls. *p<0.05, and ****p<0.0001.

### HuR is highly expressed in orthotopic PDAC fat tissue

Following our observation that adipose tissue anabolism is impaired in PDAC animals, we next sought to understand the mechanism of this phenomenon. We used Ingenuity Pathway Analysis upstream regulator analysis to identify potential candidates that could drive impaired anabolism in PDAC adipose tissue. From the DEGs identified in gWAT of PDAC mice, we found that canonical regulators of adipogenesis, such as troglitazone, fenofibrate, PPARA, and PGC1A, were predicted to be inhibited. Inflammatory cytokines associated with PDAC and cachexia, such as MYD88, IL-6, TNFA, IFNG, and IL-1B, were predicted to be activated. Among other targets that were predicted to be activated but were not typically associated with cachexia was human antigen R (HuR, *ELAVL1*), an RNA-binding protein recently established as a suppressor of adipogenesis [27] (**Figure 6A**). Given the predicted activation of HuR in gWAT and existing literature, we next asked if HuR was more abundant in adipose tissue from PDAC mice. We performed immunohistochemical staining of HuR in formalin-fixed, paraffin-embedded sham and PDAC gWAT, iWAT, pancreas, and muscle tissue. HuR staining was significantly increased in the pancreas, gWAT, and iWAT of PDAC mice versus sham controls (**Figure 6B-D, S7-8).** We also observed increased HuR protein abundance by western blotting whole gWAT tissue (**Figure 6E, S4**).

**Figure 6:**
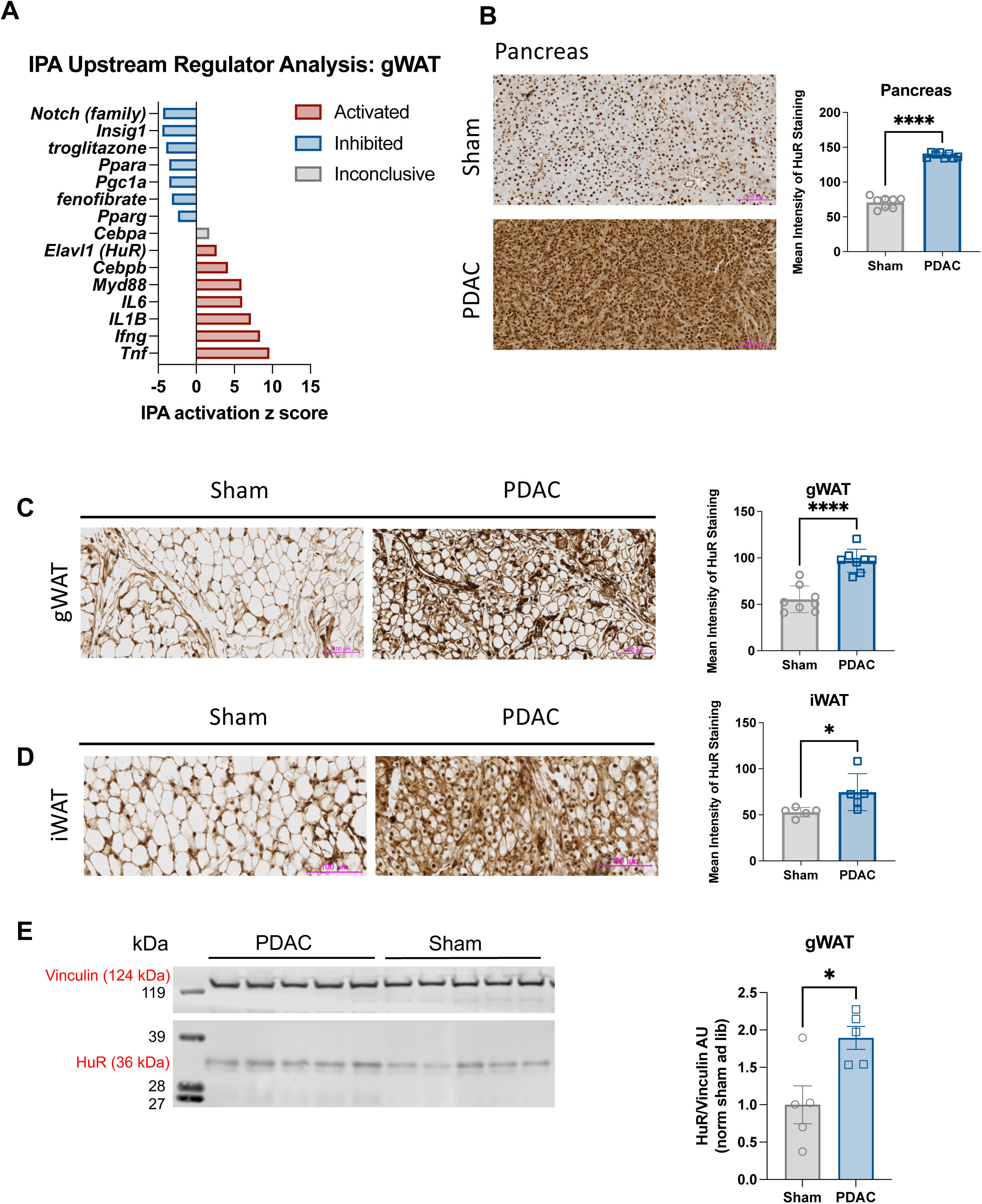
HuR is highly expressed in PDAC orthotopic pancreas and fat tissue. **(A)** Selected upstream regulator predictions from Ingenuity Pathway Analysis of differentially expressed genes in gWAT. **(B-D)** Representative images and quantification of immunohistochemical detection of HuR in tissue from PDAC and sham mice. **(B)** Pancreas N=4 male and 4 female mice per group. **(C)** gWAT N=4 male and 4 female mice per group. **(D)** iWAT N=3 female, 2 male sham and 2 female, 3 male PDAC. (**E**) Western blot analysis of PDAC and Sham gWAT protein for HuR. N=5 male mice per group. HuR protein (36 kDa) normalized to vinculin protein (124 kDa). Scale bars represent 100 um. Statistically tested with t-test. *p<0.05, **p<0.01, ***p<0.001, and ****p<0.0001.

### HuR inhibition improves adipose anabolism after fasting in PDAC mice

Since HuR has been established to drive pro-survival pathways in PDAC, we next wanted to determine if preventing HuR binding to target mRNAs could reverse adipose wasting in PDAC cachexia [26]. To do this, we treated PDAC mice with a selective HuR inhibitor, KH-3, which prevents binding of HuR to its target mRNA sequence [33]. All mice received orthotopic PDAC tumor injections and were fed *ad libitum* then fasted 24h with or without a 24h refeed at mid-cachexia (9 days after injection). In each feeding group, mice were treated with the HuR inhibitor KH-3 (100 mg/kg) or vehicle at 6-, 8-, and 10-days post-injection. Cumulative food intake between vehicle and KH-3 groups was not statistically significant prior to fast. KH-3 treated PDAC mice gained significantly more weight during refeeding compared to vehicle-treated mice (**Figure 7A-B**). Refeeding was associated with larger tumors in both vehicle and KH-3 groups, likely due to both log growth and the propensity for mice to become dehydrated during fasting (**Figure 7C**). KH-3 treatment itself had no effect on tumor growth over this time interval. In the presence of KH-3, refeeding increased WAT mass of PDAC animals, which was not observed in vehicle-treated PDAC mice (**Figure 7D-F**). These data indicate that mice treated with KH-3 shows signs of functional anabolism that are lacking in PDAC mice given vehicle treatment. This effect is evident despite the fact that KH-3 treated mice tended to lose more WAT mass during fasting than vehicle-treated mice. Pro-anabolic effects of KH-3 are specific to adipose tissue, as gastrocnemius muscle mass was not changed between fast/refeed, or vehicle/KH-3 treatment (**Figure 7F**). KH-3 treatment did not restore expression of adipogenic (*Cebpa, Adipoq, Lep*, *Retn*) or lipogenic (*Fasn, Dgat1, Dgat2, Nr1h3, Scd1, Srebp1c*) genes (**Figure 8B-C, E-F**). HuR inhibition also did not impact lipolysis gene expression (*Atgl* and *Lipe*) (**Figure 8A, D**). Thus, we show a role for HuR in adipose tissue metabolism during PDAC-associated cachexia, whereby inhibiting HuR improved anabolism and adipose retention, although without restoring the expression of genes associated with adipogenesis and lipogenesis.

**Figure 7:**
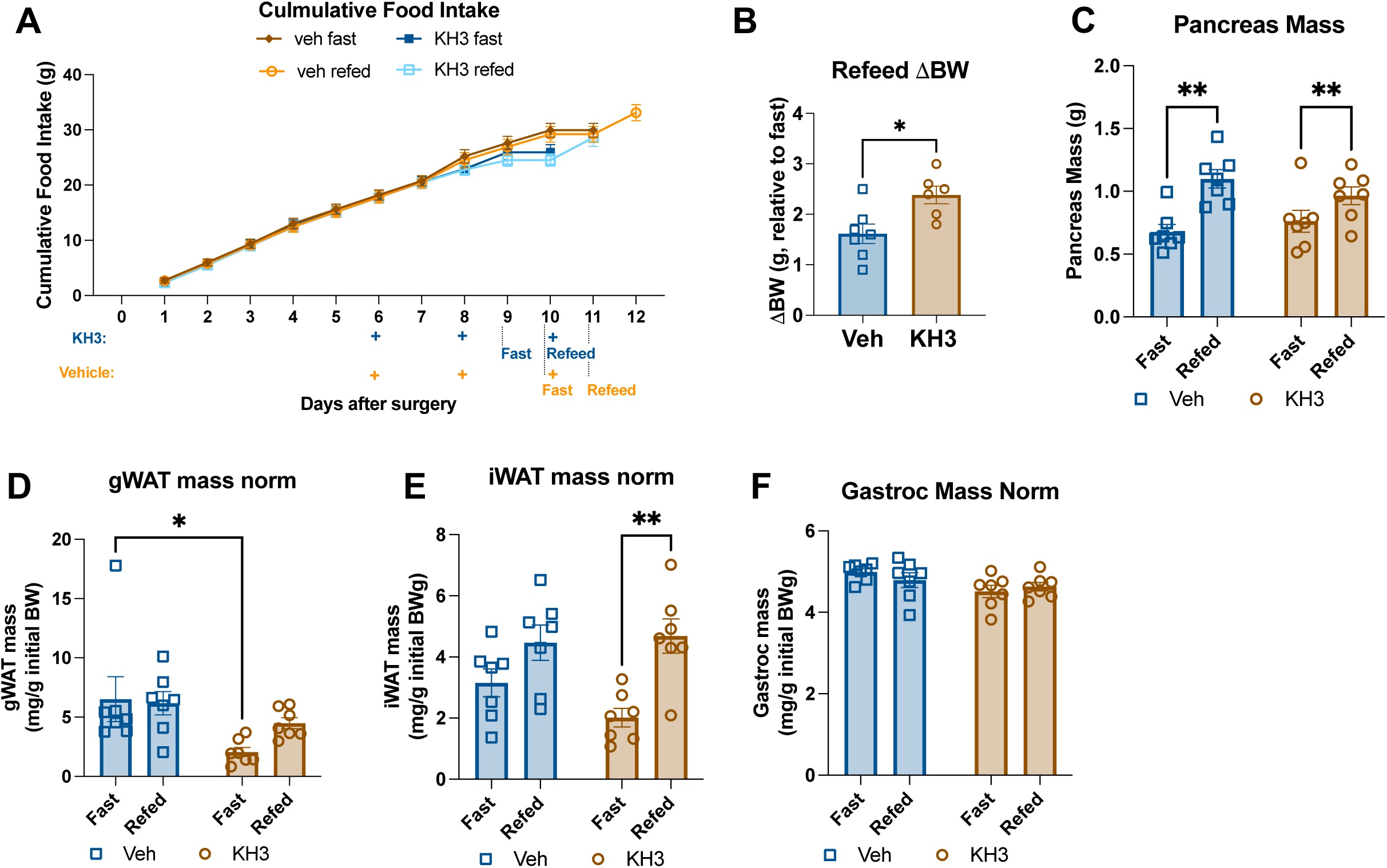
HuR inhibition improves adipose anabolism after fasting in PDAC mice. Wildtype C57BL/6J mice with PDAC were fed *ad libitum* for 9 days then fasted 24 h with or without a 24 h refeed. In each feeding paradigm, groups were treated with either vehicle or the HuR antagonist, KH-3, then euthanized 10 or 11 days after tumor implantation following the 24 h fast or 24 h refeed. Vehicle-treated mice were pair-fed to the KH-3 treated group’s average food intake from 8 days post-implantation until study end. N=7 male mice per group. **(A)** Cumulative food intake, statistically tested as an unpaired t-test of vehicle vs KH-3 treatment at day 9 (prior to fast) p =0.1121. **(B)** Change in body weight after refeeding, statistically tested with unpaired t-test. **(C)** Terminal pancreas mass. **(D)** Terminal gWAT mass. **(E)** Terminal iWAT mass. **(F)** Terminal gastrocnemius mass. 2x2 analyses were statistically tested with two-way ANOVA with Tukey multiple comparisons*p<0.05, and **p<0.01.

**Figure 8:**
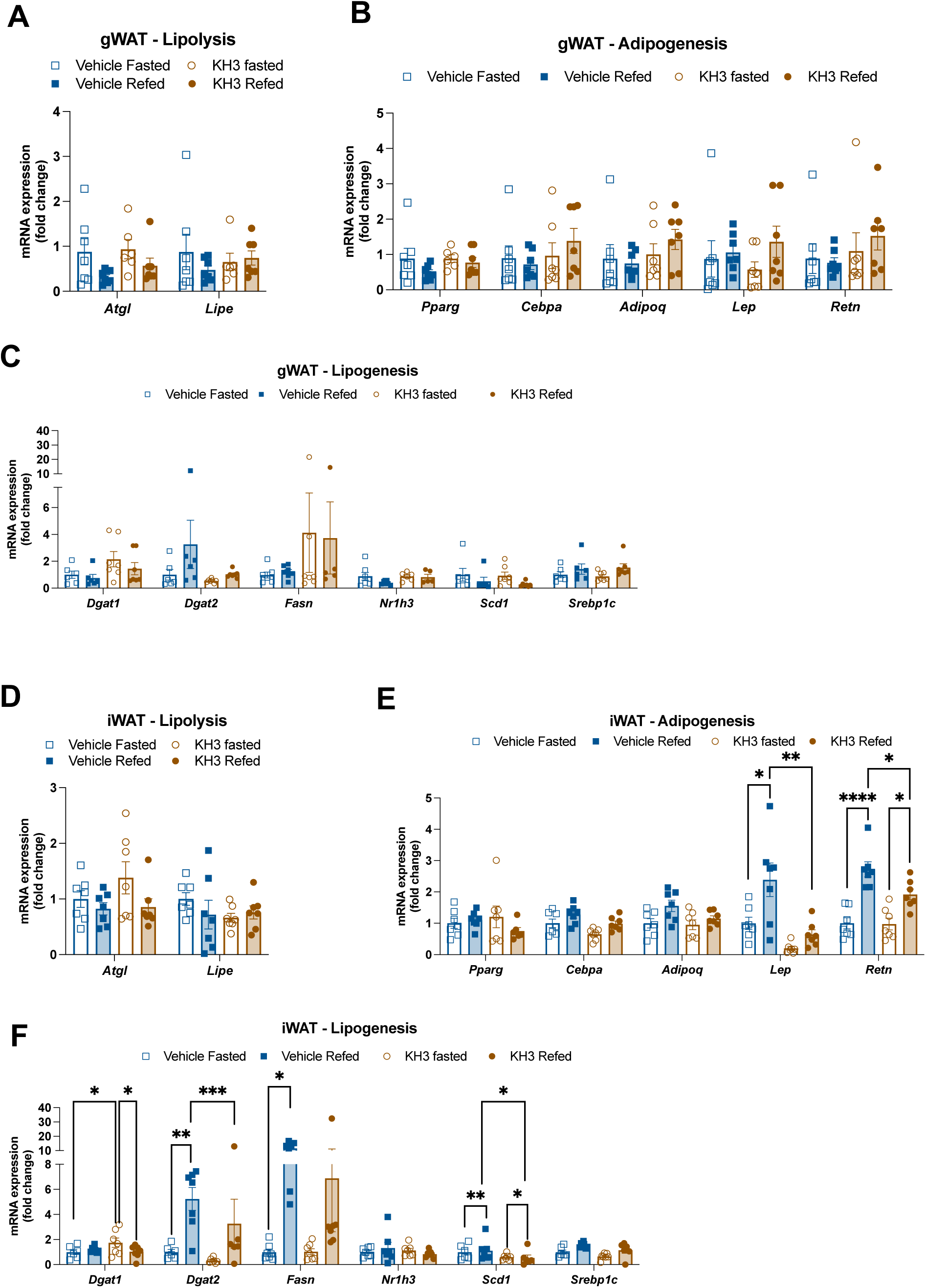
HuR inhibition does not improve gene expression patterns associated with anabolism. mRNA levels of lipolysis. **(A),** adipogenesis **(B),** and lipogenesis **(C)** genes in gWAT, normalized to beta actin expression. mRNA levels of lipolysis **(D),** adipogenesis **(E),** and lipogenesis **(F)** genes in iWAT, normalized to beta actin expression. Data were statistically tested with two-way ANOVA with Tukey multiple comparisons correction. N=7 male mice per group. *p<0.05, **p<0.01, ***p<0.001, and ****p<0.0001.

## DISCUSSION

This work evaluates the relative contributions of enhanced catabolism and impaired anabolism on fat wasting by investigating adipose tissue response to different nutritional contexts and HuR inhibition in cachectic mice. We demonstrate that our murine model of PDAC cachexia exhibits a near-complete deficit in adipose tissue anabolism in the context of reduced lipolysis. We further identify HuR as a potential molecular mediator of adipogenesis during pancreatic cancer progression. In cultured adipocytes, we showed that PDAC conditioned media does not induce a strong lipolytic response. Although the sensitivity of the glycerol release assay was limited by the absence of BSA as a free fatty acid acceptor, lipolysis and lipogenesis protein and mRNA expression data were congruent with later findings in vivo. Altogether, this emphasizes that PDAC-derived factors are sufficient to impair adipose anabolism, which corroborates a growing body of literature that challenges the dogma that fat wasting occurs as a result of only enhanced catabolism [16–18].

Our study implicates the RNA-binding protein HuR in the inhibition of adipogenesis and lipogenesis in adipose tissue. HuR can impact the transcriptome through both RNA binding and HuR-dependent splicing [26]. One documented mechanism of HuR-mediated adipogenesis regulation is via HuR binding to adipogenesis upstream regulator *Pparg*, which suppresses *Adipoq* protein expression in a non-transcriptionally dependent manner [34]. In our RNA sequencing, we did not observe increased HuR expression, although others reported increased HuR expression in response to nuclear factor-kappa B (NF-κB) signaling [35].

Alternatively, we identified HuR as an upstream regulator based on adipose transcriptomic changes associated with PDAC. In this context, HuR is more likely controlled post-transcriptionally, although further investigation is needed to identify the specific PDAC-associated factors that may activate HuR stability and nuclear translocation [36].

Small molecule antagonists of HuR, such as KH-3, are currently in development and have been found to have anti-tumor effects [26, 37]. In our studies, KH-3 restored adipose anabolism but also induced severe hemolytic anemia in all treated mice (**Figure S9**). Tumor size was not impacted by KH-3 treatment, likely because of the short period of dosing. However, studies aimed to evaluate tumor growth over 5 weeks documented slower tumor growth and fewer metastases in KH-3-treated mice [38]. KH-3 did not reverse the transcriptional repression of adipogenesis and lipogenesis observed in WAT from PDAC mice. If KH-3 restores adipogenesis by blocking HuR binding to RNA, we interpret these data to suggest that HuR acts post-transcriptionally to suppress adipose anabolism, although we cannot rule out non-HuR-mediated (i.e., off-target) mechanisms of action. In our studies, the anti-cachectic effects of KH-3 were specific to adipose tissue and did not preserve skeletal muscle mass. Although prior studies link muscle and adipose wasting, these models are all characterized by enhanced lipolysis and browning [30]. As evidenced by the adipose-specific benefits of KH-3 with concurrent muscle wasting, in our model, adipose and muscle wasting appear to be driven by independent mechanisms. Continued improvement of HuR antagonists could lead to therapeutics that restore anabolic potential in adipose tissue, providing a novel approach to alleviating one symptom of cachexia, while also providing anti-tumor benefits.

In addition to impaired anabolic potential in adipose tissue, our PDAC model also presents suppressed lipolysis. Suppressed lipolysis could contribute to systemic defects in energy utilization in cancer cachexia. Prior work from our group shows that PDAC impairs hepatic lipid oxidation but that this does not result in lipid accumulation or fatty liver [32]. It is possible that impaired lipolysis contributes to global metabolic disruption by limiting circulating lipid substrates relative to the degree of caloric deficit. Further interrogation is needed to understand how improved fat retention through HuR inhibition might impact lipid mobilization and systemic metabolism.

Our work identifies impaired adipose anabolism and increased HuR signaling as unique components of adipose wasting in cachexia. Although much of the focus in cachexia research and treatment is on skeletal muscle, multiple recent reports demonstrate that adipose loss in PDAC cachexia is a poor prognostic indicator, independent of muscle loss [5, S9-10]. This further supports existing work describing heterogeneity within cancer cachexia patients [39].

Moving forward, accounting for cachexia patient heterogeneity, including adipose wasting and loss of anabolic adipose potential, in clinical trial design may lead to increased therapeutic success. The work presented here provides a strong foundation for continuing to evaluate therapeutics designed to improve adipogenesis and lipogenesis in the context of PDAC cachexia.

## ACKNOWLEDGEMENTS

We thank all members of the Aaron Grossberg and Jonathan Brody labs for their helpful discussion and suggestions. We also would like to thank Dr. Laing Xu (Department of Molecular Biosciences, University of Kansas, Lawrence, Kansas) for generously providing us with the KH-3 compound. Author contributions are: Conceptualization, KP, AJG. Methodology, KP, HM, PP, GM, AC, AJG. Validation, PCAW, KP, HM, BW, PP. Formal Analysis, PCAW, KP, HM, BW, PP, GM, AC, AJG. Investigation, PCAW, KP, HM, BW, PP, GM, AC. Writing—Original Draft, PCAW, BLW. Writing – Review and Editing, PCAW, PP, JB, AJG. Visualization, PCAW, KP, HM, BW, PP, GM, AC. Supervision, JB, AJG. Project Administration, JB, AJG. Funding Acquisition, JB, AJG. All authors approved this manuscript.

## Ethical standards

The authors of this manuscript certify that they comply with the ethical guidelines for authorship and publishing in the Journal of Cachexia, Sarcopenia and Muscle. [40] All human and animal studies were approved by the appropriate ethics committees and were therefore performed in accordance with the ethical standards laid down in the 1964 Declaration of Helsinki and its later amendments. All human subjects provided informed consent and any identifying information of individual patients has been omitted.

## REFERENCES

1. Baracos VE, Martin L, Korc M, Guttridge DC, Fearon KCH. Cancer-associated cachexia. Nat Rev Dis Primers. 2018;4:17105. doi:10.1038/nrdp.2017.105

2. von Haehling S, Anker MS, Anker SD. Prevalence and clinical impact of cachexia in chronic illness in Europe, USA, and Japan: facts and numbers update 2016. J Cachexia Sarcopenia Muscle. 2016;7:507–9. doi:10.1002/jcsm.12167

3. Fearon K, Strasser F, Anker SD, Bosaeus I, Bruera E, Fainsinger RL, et al. Definition and classification of cancer cachexia: an international consensus. Lancet Oncol. 2011;12:489–95. doi:10.1016/S1470-2045(10)70218-7

4. Fearon K, Arends J, Baracos V. Understanding the mechanisms and treatment options in cancer cachexia. Nat Rev Clin Oncol. 2013;10:90–9. doi:10.1038/nrclinonc.2012.209

5. Kays JK, Shahda S, Stanley M, Bell TM, O’Neill BH, Kohli MD, et al. Three cachexia phenotypes and the impact of fat-only loss on survival in FOLFIRINOX therapy for pancreatic cancer. Journal of cachexia, sarcopenia and muscle. 2018;9:673–84.

6. Mantovani G, Maccio A, Lai P, Massa E, Ghiani M, Santona MC. Cytokine activity in cancer-related anorexia/cachexia: role of megestrol acetate and medroxyprogesterone acetate. Semin Oncol. 1998;25:45–52.

7. Straub RH, Cutolo M, Buttgereit F, Pongratz G. Energy regulation and neuroendocrine-immune control in chronic inflammatory diseases. J Intern Med. 2010;267:543–60. doi:10.1111/j.1365-2796.2010.02218.x

8. Argiles JM, Busquets S, Toledo M, Lopez-Soriano FJ. The role of cytokines in cancer cachexia. Curr Opin Support Palliat Care. 2009;3:263–8. doi:10.1097/SPC.0b013e3283311d09

9. MacDonald N, Easson AM, Mazurak VC, Dunn GP, Baracos VE. Understanding and managing cancer cachexia. J Am Coll Surg. 2003;197:143-61. doi:10.1016/S1072-7515(03)00382-X

10. Carson JL, Hernandez M, Jaspers I, Mills K, Brighton L, Zhou H, et al. Interleukin-13 stimulates production of nitric oxide in cultured human nasal epithelium. In Vitro Cell Dev Biol Anim. 2018;54:200–4. doi:10.1007/s11626-018-0233-y

11. Haslett PA. Anticytokine approaches to the treatment of anorexia and cachexia. Semin Oncol. 1998;25:53–7.

12. Moldawer LL, Copeland EM, 3rd. Proinflammatory cytokines, nutritional support, and the cachexia syndrome: interactions and therapeutic options. Cancer. 1997;79:1828–39.

13. Das SK, Eder S, Schauer S, Diwoky C, Temmel H, Guertl B, et al. Adipose triglyceride lipase contributes to cancer-associated cachexia. Science. 2011;333:233–8.

14. Petruzzelli M, Schweiger M, Schreiber R, Campos-Olivas R, Tsoli M, Allen J, et al. A switch from white to brown fat increases energy expenditure in cancer-associated cachexia. Cell metabolism. 2014;20:433–47.

15. Taylor J, Uhl L, Moll I, Hasan SS, Wiedmann L, Morgenstern J, et al. Endothelial Notch1 signaling in white adipose tissue promotes cancer cachexia. Nature Cancer. 2023;4:1544–60.

16. Langer HT, Ramsamooj S, Dantas E, Murthy A, Ahmed M, Ahmed T, et al. Restoring adiponectin via rosiglitazone ameliorates tissue wasting in mice with lung cancer. Acta Physiologica. 2024;e14167.

17. Jang HJ, Kim HY, Lyu JH, Muthamil S, Shin UC, Kim HS, et al. Bee (Apis mellifera L. 1758) wax restores adipogenesis and lipid accumulation of 3T3-L1 cells in cancer-associated cachexia condition. Food Science & Nutrition. 2024;

18. Tien S-C, Chang C-C, Huang C-H, Peng H-Y, Chang Y-T, Chang M-C, et al. Exosomal miRNA 16-5p/29a-3p from pancreatic cancer induce adipose atrophy by inhibiting adipogenesis and promoting lipolysis. Iscience. 2024;27:

19. Cook KB, Kazan H, Zuberi K, Morris Q, Hughes TR. RBPDB: a database of RNA-binding specificities. Nucleic Acids Res. 2011;39:D301–8. doi:10.1093/nar/gkq1069

20. Gerstberger S, Hafner M, Tuschl T. A census of human RNA-binding proteins. Nat Rev Genet. 2014;15:829–45. doi:10.1038/nrg3813

21. Pihlajamaki J, Lerin C, Itkonen P, Boes T, Floss T, Schroeder J, et al. Expression of the splicing factor gene SFRS10 is reduced in human obesity and contributes to enhanced lipogenesis. Cell Metab. 2011;14:208–18. doi:10.1016/j.cmet.2011.06.007

22. Brosch M, von Schonfels W, Ahrens M, Nothnagel M, Krawczak M, Laudes M, et al. SFRS10--a splicing factor gene reduced in human obesity? Cell Metab. 2012;15:265–6; author reply 7-9. doi:10.1016/j.cmet.2012.02.002

23. Huot ME, Vogel G, Zabarauskas A, Ngo CT, Coulombe-Huntington J, Majewski J, et al. The Sam68 STAR RNA-binding protein regulates mTOR alternative splicing during adipogenesis. Mol Cell. 2012;46:187–99. doi:10.1016/j.molcel.2012.02.007

24. Chou CF, Lin YY, Wang HK, Zhu X, Giovarelli M, Briata P, et al. KSRP ablation enhances brown fat gene program in white adipose tissue through reduced miR-150 expression. Diabetes. 2014;63:2949–61. doi:10.2337/db13-1901

25. Dai N, Zhao L, Wrighting D, Kramer D, Majithia A, Wang Y, et al. IGF2BP2/IMP2-Deficient mice resist obesity through enhanced translation of Ucp1 mRNA and Other mRNAs encoding mitochondrial proteins. Cell Metab. 2015;21:609–21. doi:10.1016/j.cmet.2015.03.006

26. Finan JM, Sutton TL, Dixon DA, Brody JR. Targeting the RNA-binding protein HuR in cancer. Cancer Research. 2023;83:3507–16.

27. Siang DTC, Lim YC, Kyaw AMM, Win KN, Chia SY, Degirmenci U, et al. The RNA-binding protein HuR is a negative regulator in adipogenesis. Nat Commun. 2020;11:213. doi:10.1038/s41467-019-14001-8

28. Michaelis KA, Zhu X, Burfeind KG, Krasnow SM, Levasseur PR, Morgan TK, et al. Establishment and characterization of a novel murine model of pancreatic cancer cachexia. J Cachexia Sarcopenia Muscle. 2017;8:824–38. doi:10.1002/jcsm.12225

29. Schweiger M, Eichmann TO, Taschler U, Zimmermann R, Zechner R, Lass A. Measurement of lipolysis. In: Elsevier; 2014. pp. 171-93.

30. Rupert JE, Narasimhan A, Jengelley DH, Jiang Y, Liu J, Au E, et al. Tumor-derived IL-6 and trans-signaling among tumor, fat, and muscle mediate pancreatic cancer cachexia. Journal of Experimental Medicine. 2021;218:e20190450.

31. Liberzon A, Birger C, Thorvaldsdóttir H, Ghandi M, Mesirov JP, Tamayo P. The molecular signatures database hallmark gene set collection. Cell systems. 2015;1:417–25.

32. Arneson-Wissink PC, Mendez H, Pelz K, Dickie J, Bartlett AQ, Worley BL, et al. Hepatic signal transducer and activator of transcription-3 signalling drives early-stage pancreatic cancer cachexia via suppressed ketogenesis. J Cachexia Sarcopenia Muscle. 2024;15:975–88. doi:10.1002/jcsm.13466

33. Wu X, Gardashova G, Lan L, Han S, Zhong C, Marquez RT, et al. Targeting the interaction between RNA-binding protein HuR and FOXQ1 suppresses breast cancer invasion and metastasis. Commun Biol. 2020;3:193. doi:10.1038/s42003-020-0933-1

34. Hwang JS, Lee WJ, Hur J, Lee HG, Kim E, Lee GH, et al. Rosiglitazone-dependent dissociation of HuR from PPAR-γ regulates adiponectin expression at the posttranscriptional level. The FASEB Journal. 2019;33:7707–20. doi:10.1096/fj.201802643r

35. Kang MJ, Ryu BK, Lee MG, Han J, Lee JH, Ha TK, et al. NF-&#x3ba;B Activates Transcription of the RNA-Binding Factor HuR, via PI3K-AKT Signaling, to Promote Gastric Tumorigenesis. Gastroenterology. 2008;135:2030–42.e3. doi:10.1053/j.gastro.2008.08.009

36. Grammatikakis I, Abdelmohsen K, Gorospe M. Posttranslational control of HuR function. Wiley Interdisciplinary Reviews: RNA. 2017;8:e1372.

37. Huang Z, Liu S, Tang A, Wu X, Aube J, Xu L, et al. Targeting RNA-binding protein HuR to inhibit the progression of renal tubular fibrosis. Journal of Translational Medicine. 2023;21:428.

38. Dong R, Chen P, Polireddy K, Wu X, Wang T, Ramesh R, et al. An RNA-Binding Protein, Hu-antigen R, in Pancreatic Cancer Epithelial to Mesenchymal Transition, Metastasis, and Cancer Stem Cells. Mol Cancer Ther. 2020;19:2267–77. doi:10.1158/1535-7163.Mct-19-0822

39. Fearon Kenneth CH, Glass David J, Guttridge Denis C. Cancer Cachexia: Mediators, Signaling, and Metabolic Pathways. Cell Metabolism. 2012;16:153–66. doi:10.1016/j.cmet.2012.06.011

40. von Haehling S, Coats AJ, Anker SD. Ethical guidelines for publishing in the Journal of Cachexia, Sarcopenia and Muscle: update 2021. Wiley Online Library; 2021. pp. 2259–61.

