## Supplement for "Impaired adipose anabolism in pancreatic cancer cachexia is reversed by HuR inhibition"

### SUPPLEMENTAL METHODS

*RNAsequencing Analysis*

Reference genome and gene model annotation files were downloaded from genome website browser (NCBI/UCSC/Ensembl) directly. FASTQ files obtained from sequencing were put through fastp to trim adapters and then fastqc for quality analysis. STAR 2.7.1a was then used to align to reference genome (mm10) to trimmed reads after which htseq was used to obtain gene counts. Raw reads were normalized using DESeq2 (1.38.3). Genes with an adjusted P-value <0.05 found by DESeq2 were assigned as differentially expressed. Gene set enrichment analysis (GSEA) was performed through the Broad Institute GUI. Normalized counts from DESeq2 from gWAT were input and analyzed with relationship to the Molecular Signature Database (msigdb). Significant Hallmark genes (p <0.05) were graph from both positive and negative normalized enrichment scores (NES). Log2 fold change, p-values and p-adj from differential expression data obtained through deseq2 were loaded into Qiagen's Ingenuity Pathway Analysis (IPA) to look at upstream and downstream regulators. Volcano plots created from differential expression data of pairwise comparison with sham groups being baseline using ggplot2 (3.5). Significant genes from adipogenesis pathway are labeled. Heatmaps were created of genes in the adipogenesis pathway regardless of significance using pheatmap(1.0.12).

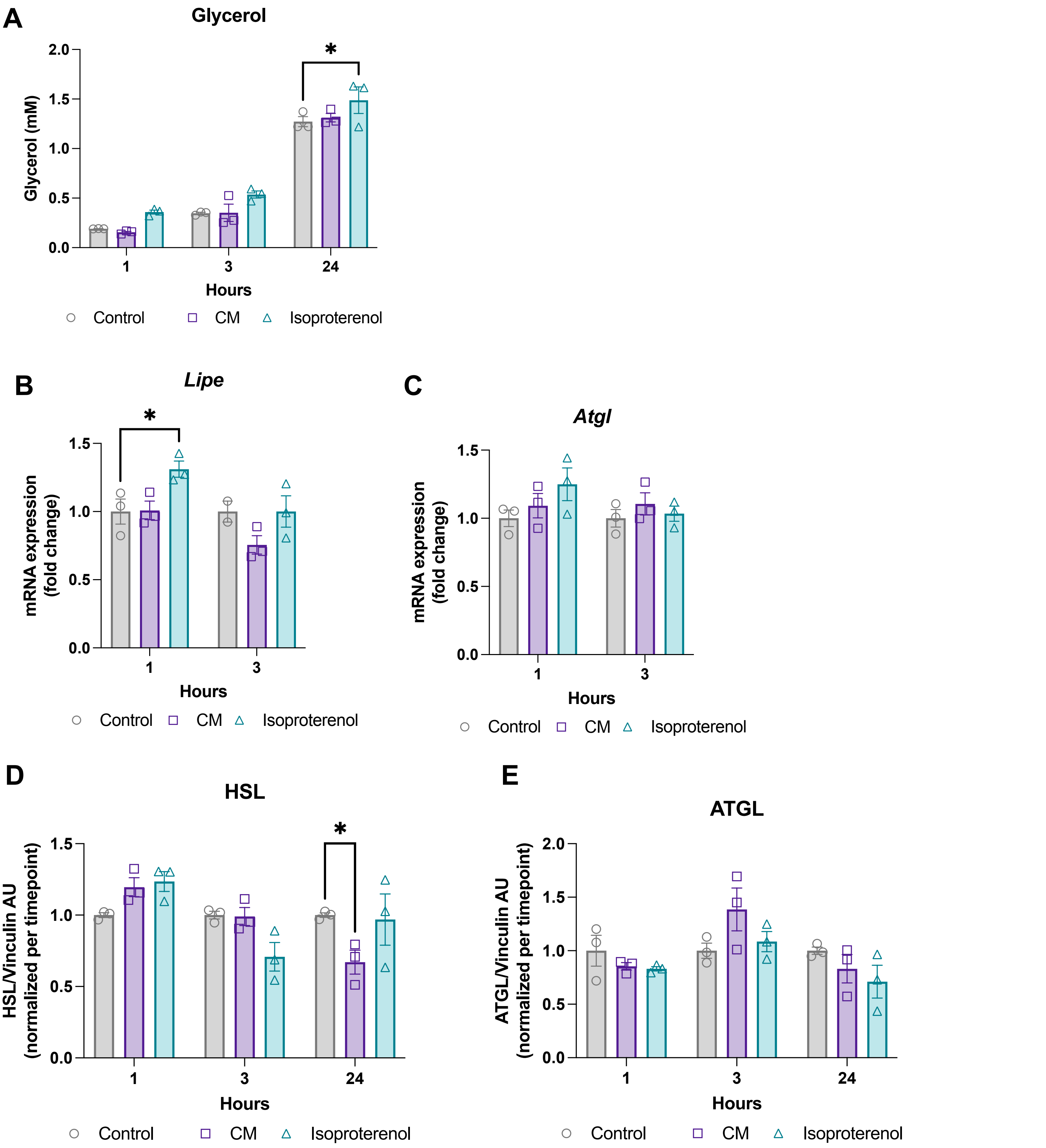


**Figure S1: Impact of KPC conditioned media on lipolysis in adipocytes.** (A) Media glycerol levels, not normalized. Conditioned media and control media alone did not contain glycerol above background (assay buffer only) levels.(B-C) mRNA levels of *Atgl* and *Lipe*. Cells collected 1 and 3 hours after KPC CM or isoproterenol (10 uM) treatment. (D-E) Western blot analysis of total HSL and ATGL protein. N=3 wells per condition. Statistically tested with two-way ANOVA with Tukey correction for multiple comparisons. $p<0.05$, $^{**}p<0.01$, $^{***}p<0.001$, and $^{****}p<0.0001$.

Figure S1

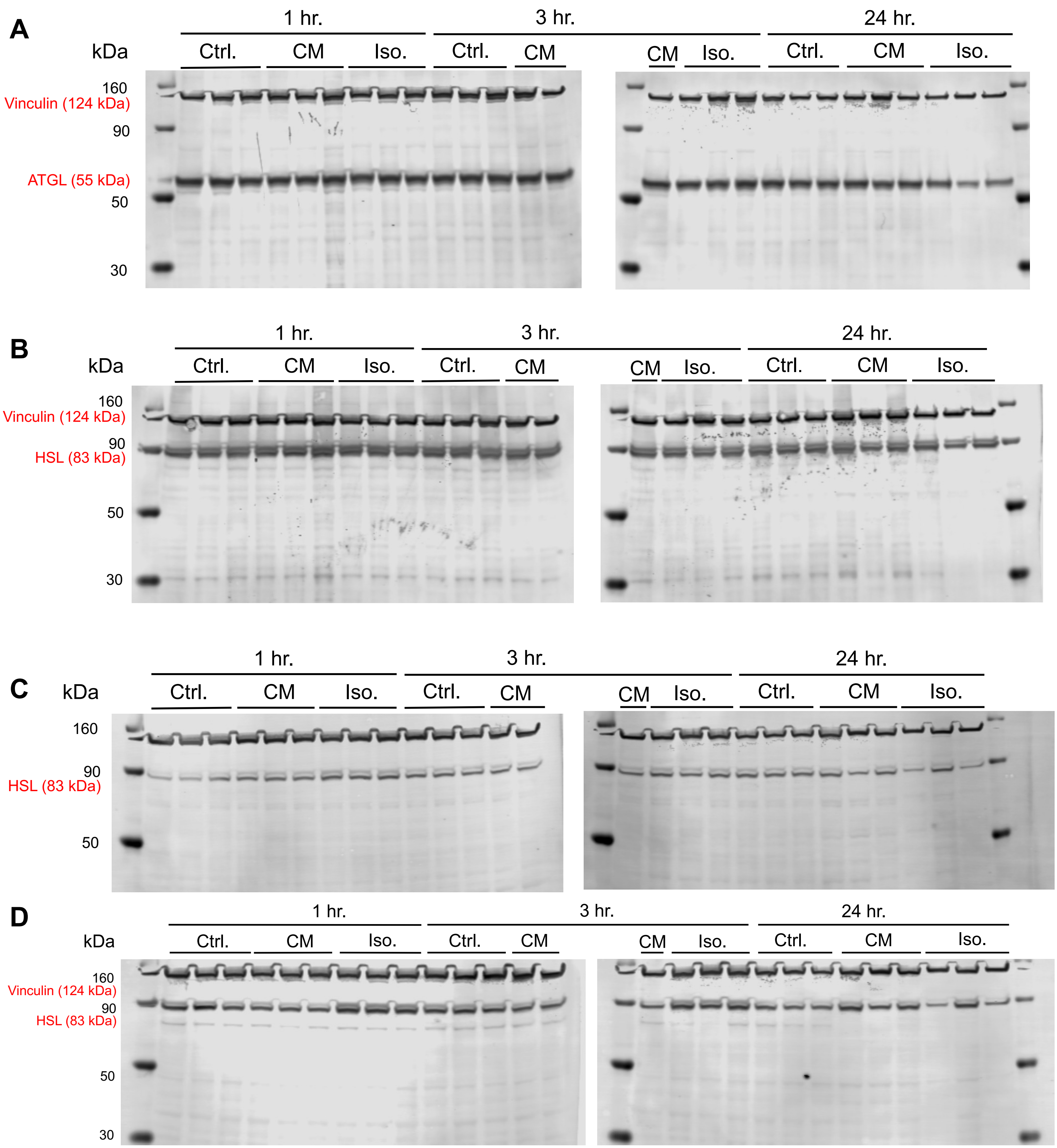


**Figure S2: 3T3-L1 protein western blot full gels.** (A) ATGL and Vinculin. (B) HSL and Vinculin. (C) p-HSL S563 and Vinculin. (D) p-HSL S660 and Vinculin. . Cells collected 1 and 3 hours after KPC CM or isoproterenol (10 uM) treatment. N=3 wells per condition.

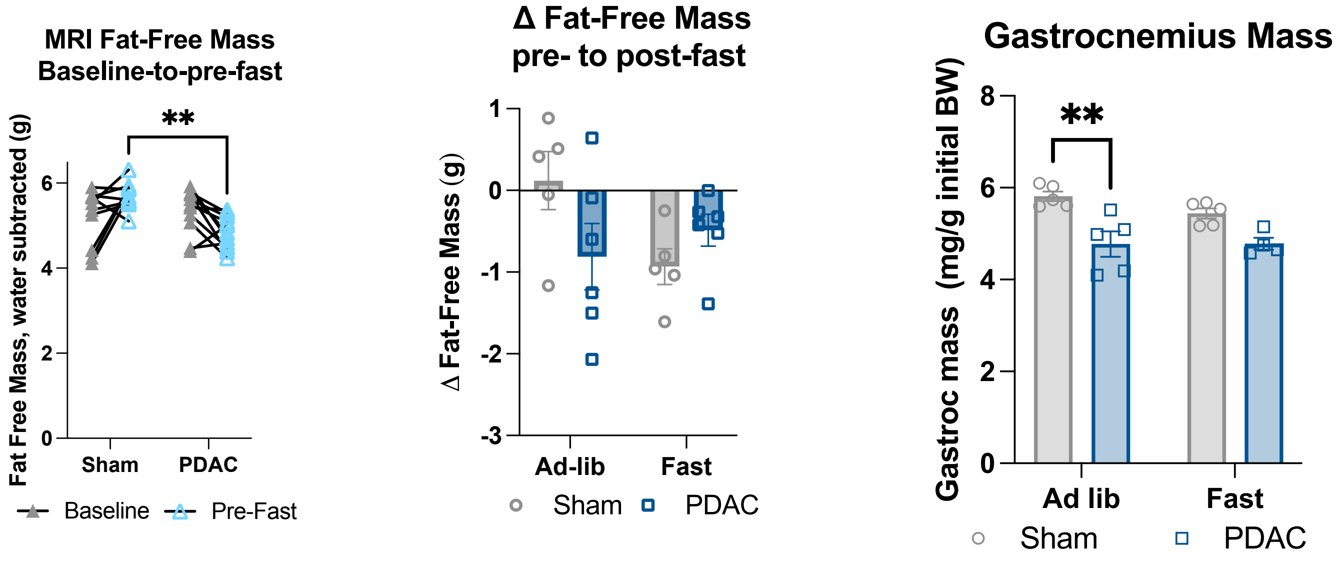


**Figure S3: Impact of fasting on lean mass in pancreatic ductal adenocarcinoma-bearing mice.** Wildtype C57BL/6J mice with PDAC or sham implantations were fed *ad libitum* or fasted 16h at mid-cachexia (9 days after injection). Animals were euthanized 10 or 11 days after tumor implantation following the 16h fast. N= 5 sham, 6 PDAC male mice per feeding condition. **(A-B)** Body composition changes in fat-free mass (water subtracted) were characterized at baseline to pre-fast between sham and PDAC mice **(A)**, and at pre- to post-fast **(B)**. **(C)** Terminal gastrocnemius muscle mass. 2x2 analyses were statistically tested with two-way ANOVA or mixed effects model with Tukey multiple comparisons. *p<0.05, **p<0.01, and ****p<0.0001.

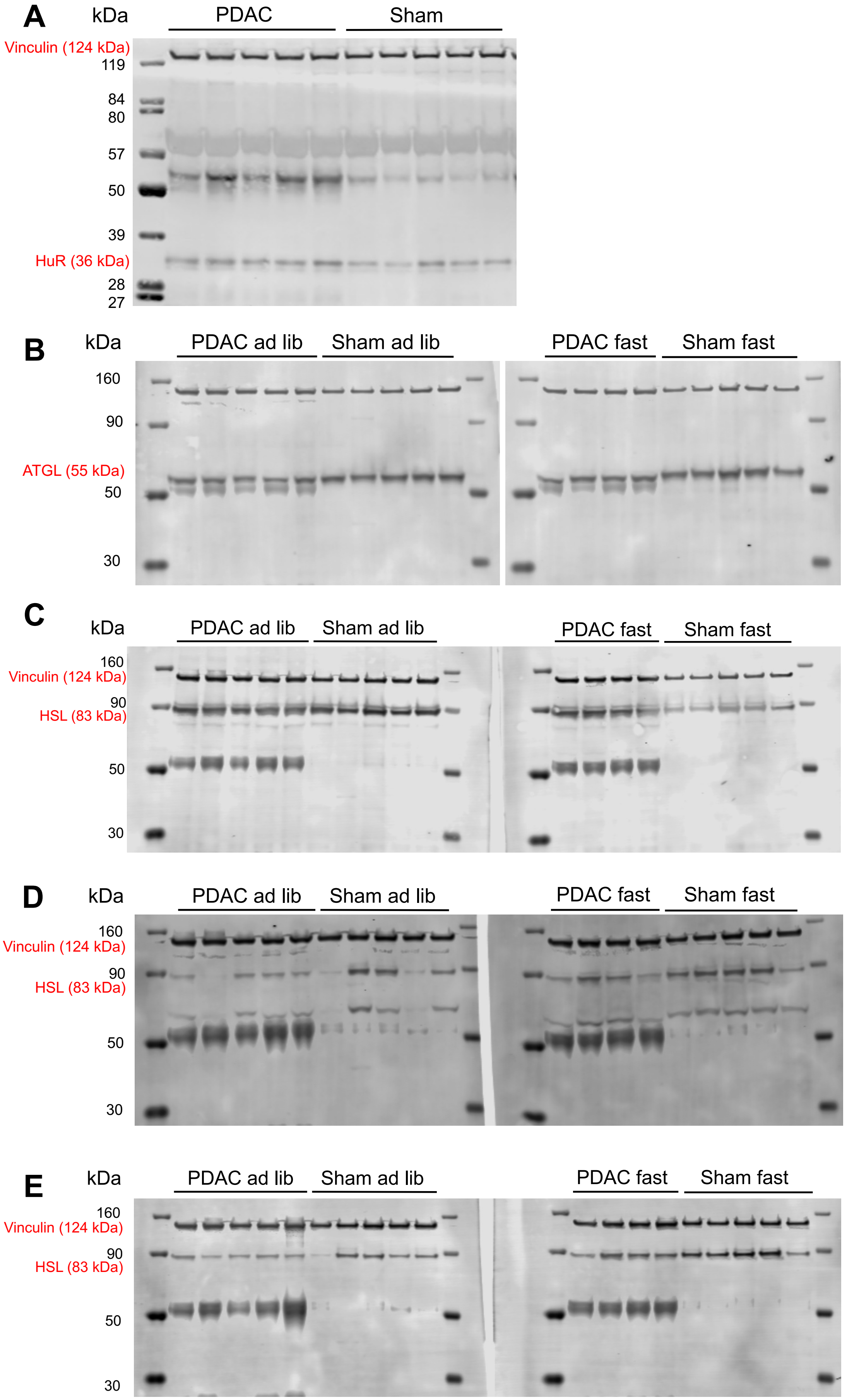


**Figure S4: gWAT protein western blot full gels.** (A) HuR and Vinculin. (B) ATGL and Vinculin. (C) HSL and Vinculin. (D) p-HSL S563 and Vinculin. (E) p-HSL S660 and Vinculin.

Figure S4

A

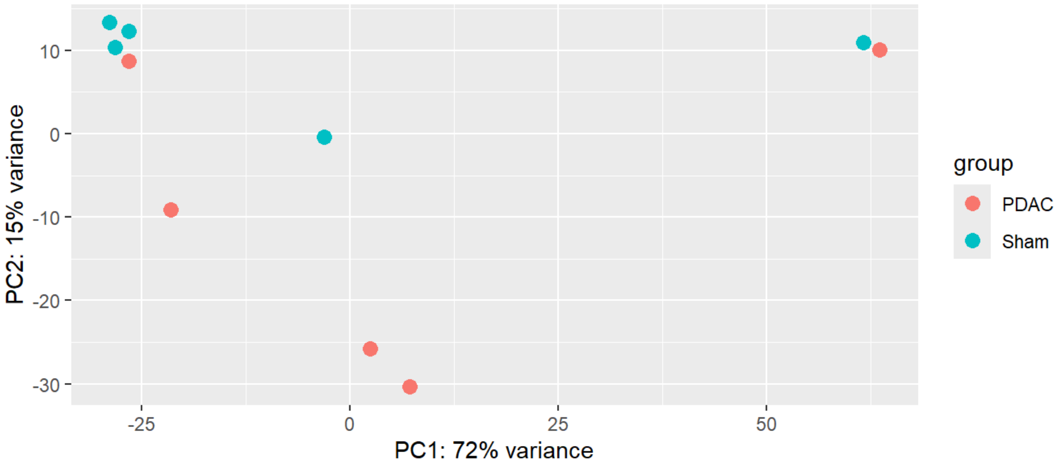


B

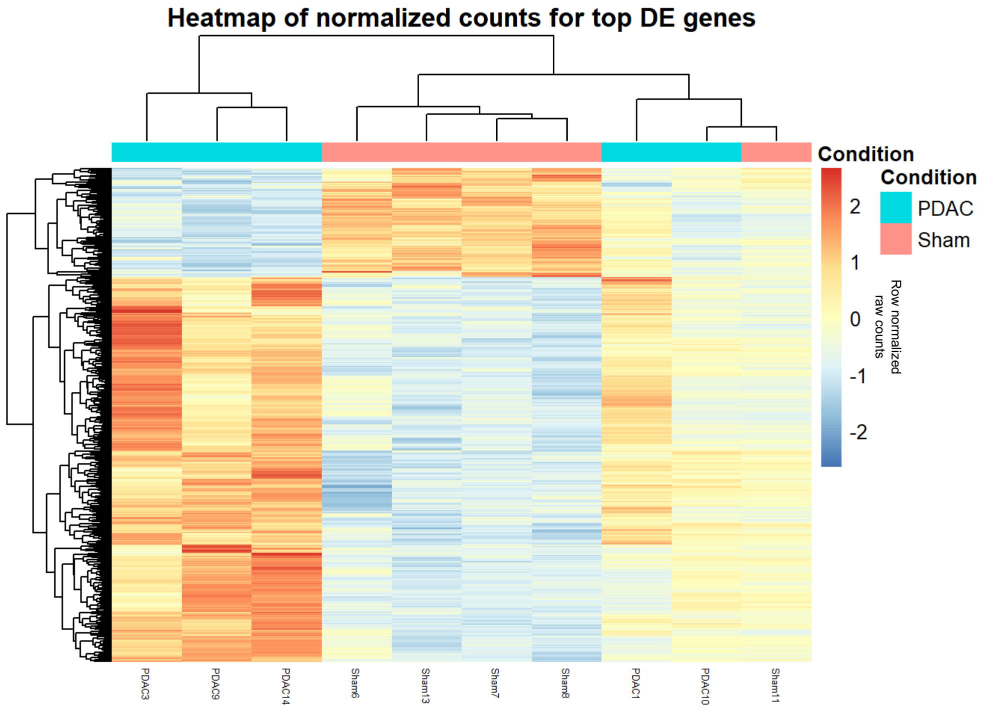


**Figure S5: Unbiased clustering of gWAT RNAseq samples. (A)** Principal component analysis of PDAC and sham iWAT samples. **(B)** Unbiased hierarchical clustering of the top differentially expressed genes.

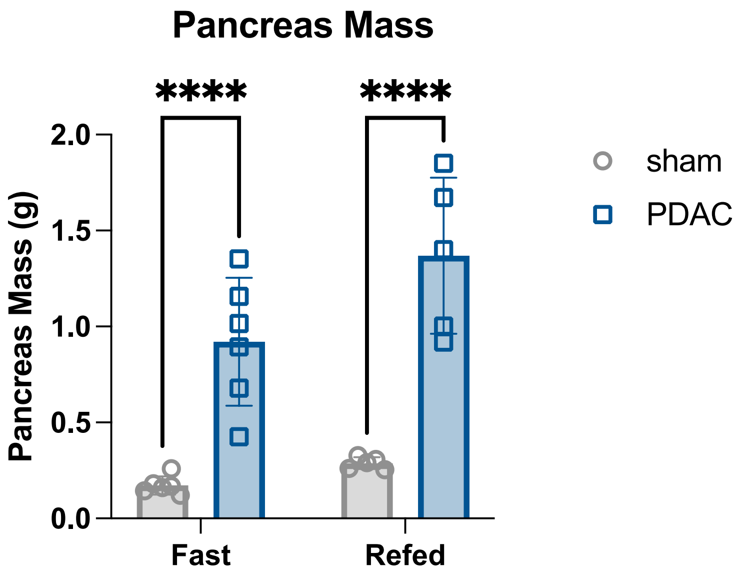


**Figure S6:** Pancreas mass (containing tumor) from sham and PDAC mice afer 24 hour fast, or 24 hour fast followed by 24 hour refeeding at 9 days post implantation. 2x2 analyses were statistically tested with two-way ANOVA or mixed effects model with Tukey multiple comparisons. *p<0.05, **p<0.01, and ****p<0.0001.

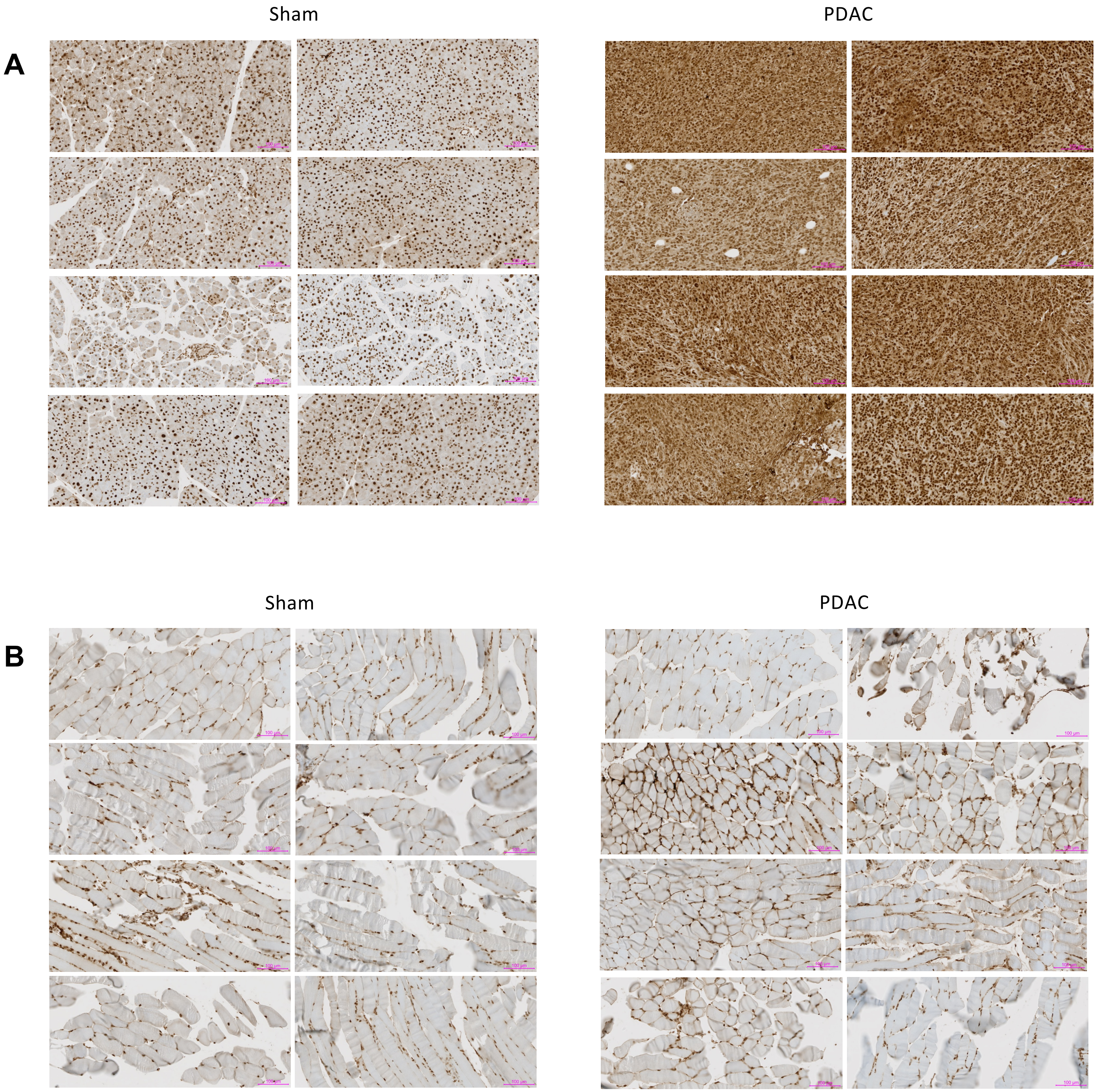


**Figure S7: Additional images of HuR staining. (A)** pancreas tissue. **(B)** gastrocnemius muscle tissue. Scale bars represent 100 um.

A

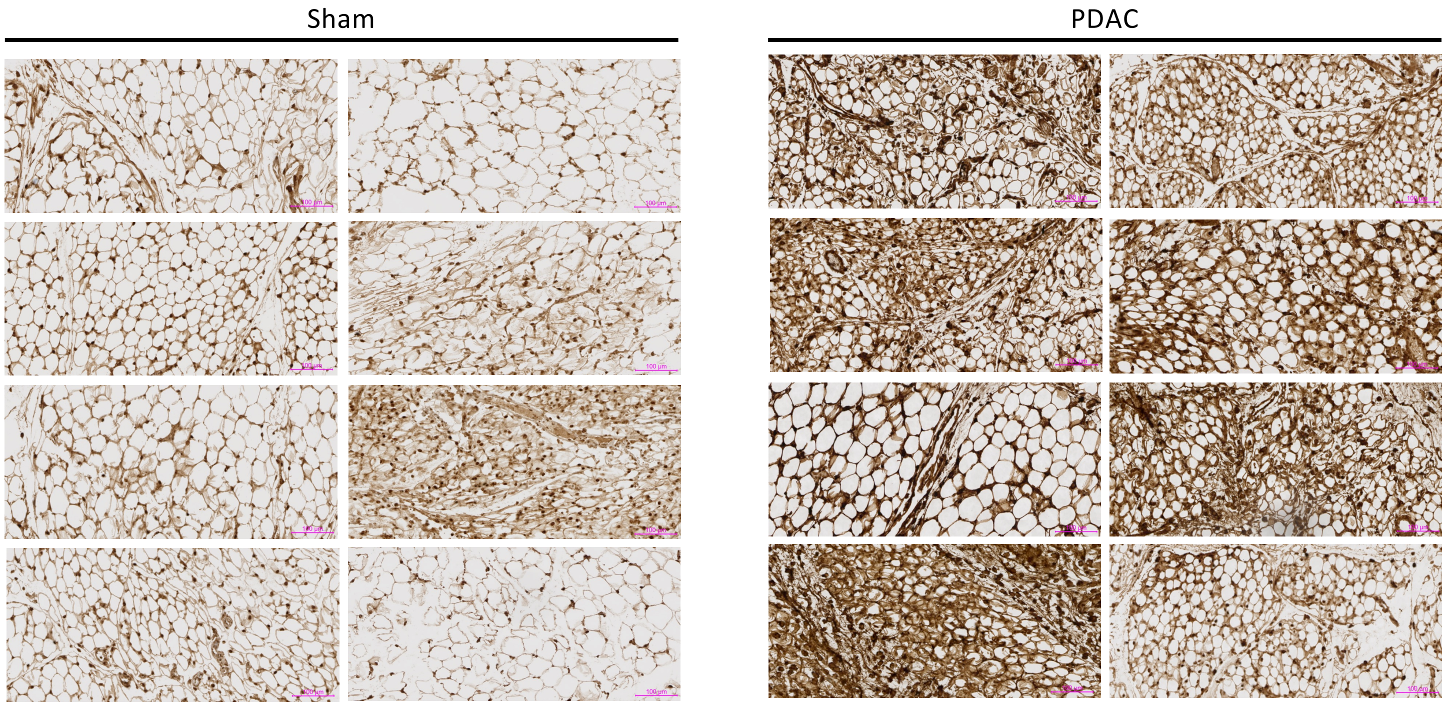


B

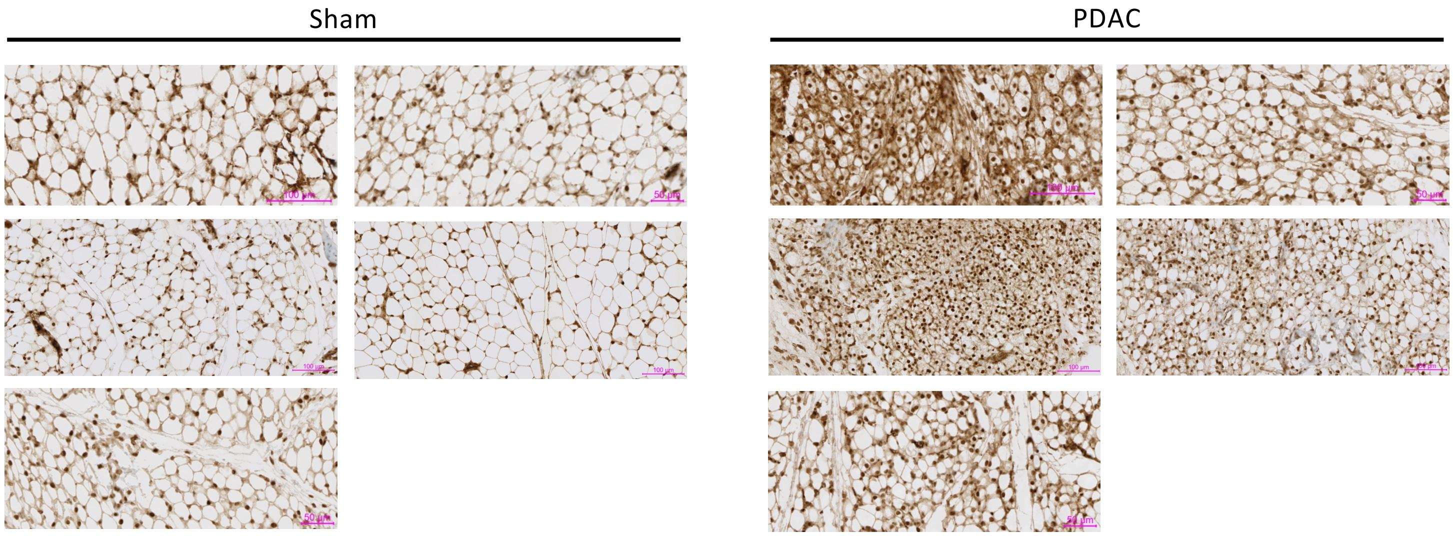


**Figure S8: Additional images of HuR staining. (A)** gWAT tissue. **(B)** iWAT tissue. Scale bars represent 100 um unless noted to be 50 um.

**A**

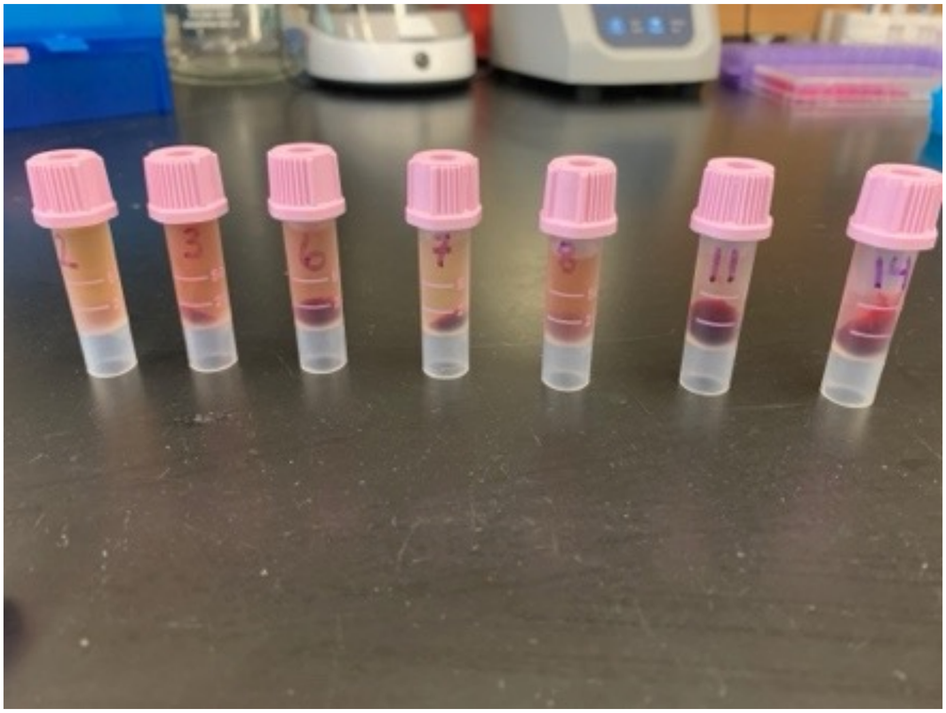

**Figure S9: Evidence of anemia in the centrifuged blood from PDAC, KH3-treated, refed mice.** EDTA-treated blood samples after centrifugation at 2,000g x 20 min to collect plasma. The red blood cell volume (hematocrit) is visibly low after centrifugation, although we did not specifically measure the Hgb or Hct of this blood.

### SUPPLEMENTAL REFERENCES


S1. Tisdale MJ. Biology of cachexia. J Natl Cancer Inst. 1997;89:1763-73. doi:10.1093/jnci/89.23.1763

S2. Grossberg AJ, Scarlett JM, Marks DL. Hypothalamic mechanisms in cachexia. Physiol Behav. 2010;100:478-89. doi:10.1016/j.physbeh.2010.03.011

S3. Olson B, Marks DL. Pretreatment Cancer-Related Cognitive Impairment-Mechanisms and Outlook. Cancers (Basel). 2019;11:doi:10.3390/cancers11050687

S4. von Haehling S, Anker SD. Cachexia as a major underestimated and unmet medical need: facts and numbers. J Cachexia Sarcopenia Muscle. 2010;1:1-5. doi:10.1007/s13539-010-0002-6

S5. Fearon KC, Glass DJ, Guttridge DC. Cancer cachexia: mediators, signaling, and metabolic pathways. Cell Metab. 2012;16:153-66. doi:10.1016/j.cmet.2012.06.011

S6. Tisdale MJ. Cachexia in cancer patients. Nat Rev Cancer. 2002;2:862-71. doi:10.1038/nrc927

S7. Aoyagi T, Terracina KP, Raza A, Matsubara H, Takabe K. Cancer cachexia, mechanism and treatment. World J Gastrointest Oncol. 2015;7:17-29. doi:10.4251/wjgo.v7.i4.17

S8. Mueller TC, Burmeister MA, Bachmann J, Martignoni ME. Cachexia and pancreatic cancer: are there treatment options? World J Gastroenterol. 2014;20:9361-73. doi:10.3748/wjg.v20.i28.9361

S9. Klassen PN, Baracos V, Ghosh S, Martin L, Sawyer MB, Mazurak VC. Muscle and adipose wasting despite disease control: unaddressed side effects of palliative chemotherapy for pancreatic cancer. Cancers. 2023;15:4368.

S10. Yu S-Y, Luan Y, Dong R, Abazarikia A, Kim S-Y. Adipose tissue wasting as a determinant of pancreatic cancer-related cachexia. Cancers. 2022;14:4754.
